# Synergy of RNA Concentration, RNA Binding Proteins, and RNA Palindrome Drives clustering of *oskar* mRNA *in vivo*

**DOI:** 10.64898/2026.04.30.721893

**Authors:** Ziqing Ye, Siran Tian, Ayse Ecer, Tatjana Trcek

## Abstract

Organization of mRNAs into clusters has been observed in many cellular contexts, yet the features that govern this process in vivo remain poorly understood. Using super-resolution microscopy, single-mRNA imaging, and genetic perturbations, we investigated how mRNA concentration, the double-stranded RNA-binding protein Staufen (Stau), and intermolecular base-pairing driven by an RNA palindrome influence clustering of *oskar* (*osk*) mRNA in *Drosophila* embryos. We find that these factors collectively optimize *osk* clustering by promoting its dimerization and subsequent oligomerization. Both processes depend on all three factors, although oligomerization is more sensitive to their perturbation, indicating that the driving force for *osk* oligomerization is partially distinct from that governing dimerization. Furthermore, expression of Stau nearly doubles the likelihood of *osk* dimerization whereas disruption of the palindrome reduces it fourfold indicating that the presence of Stau and the palindrome lowers the concentration threshold of *osk* mRNA required for dimerization. Notably, insertion of the *osk* palindrome into a reporter mRNA markedly increased its association with the endogenous *osk*, further supporting the conclusion that the palindrome potently drives intermolecular base pairing. Importantly, this experiment also identified the palindrome as the major contributor to heterotypic clustering between the endogenous *osk* and the reporter mRNA. Finally, computational analyses identified a subset of early embryonic mRNAs predicted to harbor palindromes similar to those found in *osk*. Among these, *eIF3a* mRNA emerged as a candidate whose clustering may likewise be driven by intermolecular base pairing. Together, our findings raise the possibility that mRNA clustering driven by palindrome-mediated intermolecular base pairing may be more widespread than previously appreciated and may represent an important mechanism for controlling mRNA spatial organization during early *Drosophila* development.

## Introduction

Spatial organization of messenger RNAs (mRNAs) is a widespread mechanism for controlling the spatial and temporal production of proteins within cells and is particularly impactful during organismal development, where localized mRNAs influence cell fate, morphology, and body plan formation across metazoans^1^.

Beyond being targeted to defined cellular structures such as the endoplasmic reticulum or RNA granules, mRNAs can also organize with one another to form mRNA clusters^2–12^. These clusters, which may contain only a few transcripts, arise through intermolecular interactions involving both RNA-binding proteins (RBPs) as well as the RNAs themselves^13^. In this context, RNAs can serve as scaffolds that recruit RBPs to drive mRNA clustering, or they may directly promote clustering via RNA:RNA interactions^14–19^.

Intermolecular base pairing between complementary sequences (CSs) represents a prominent mechanism of RNA-driven clustering^14,15,17,20–25^. This mechanism is well characterized in RNA viruses where genome dimerization and higher-order assembly depend on defined CS elements. For example, viruses like Influenza A, Human immunodeficiency virus-1 (HIV-1), Murine leukemia virus (MLV) and the Flock House virus (FHV) rely on intermolecular base pairing to organize and package their RNA genomes^26–34^, and disruption of these interactions blocks virion assembly^26–28,32,33^. Dimerization-competent CSs are typically structurally exposed which facilitates recognition among mRNAs and positions them for stable and persistent interactions^21,26,27,35,36^. They are also often GC-rich, which thermodynamically enhances the stability of base pairing. For example, in HIV-1, a six-nucleotide GC-rich palindrome located at the apex of a stem-loop nucleates RNA dimerization^21,37^.

Intermolecular base pairing also operates in eukaryotic mRNAs. In the filamentous fungus *Ashbya gossypii*, base pairing between *BNI1* and *SPA2* mRNAs, both essential for cell cycle progression and nuclear division, drives their organization within cells^23^. This organization is important for establishing compositional specificity of biomolecular condensates associated with polarized growth and nuclear divisions in the fungus^23^. Similarly, in *Drosophila*, complementary sequence interactions between the *oskar* (*osk*) (Fig. 1Ai,ii) and *bicoid* (*bcd*) mRNAs, two mRNAs that are critical for early embryonic development, promote their respective enrichment at distinct embryonic locations ^15,22,38^.

**FIGURE 1:**
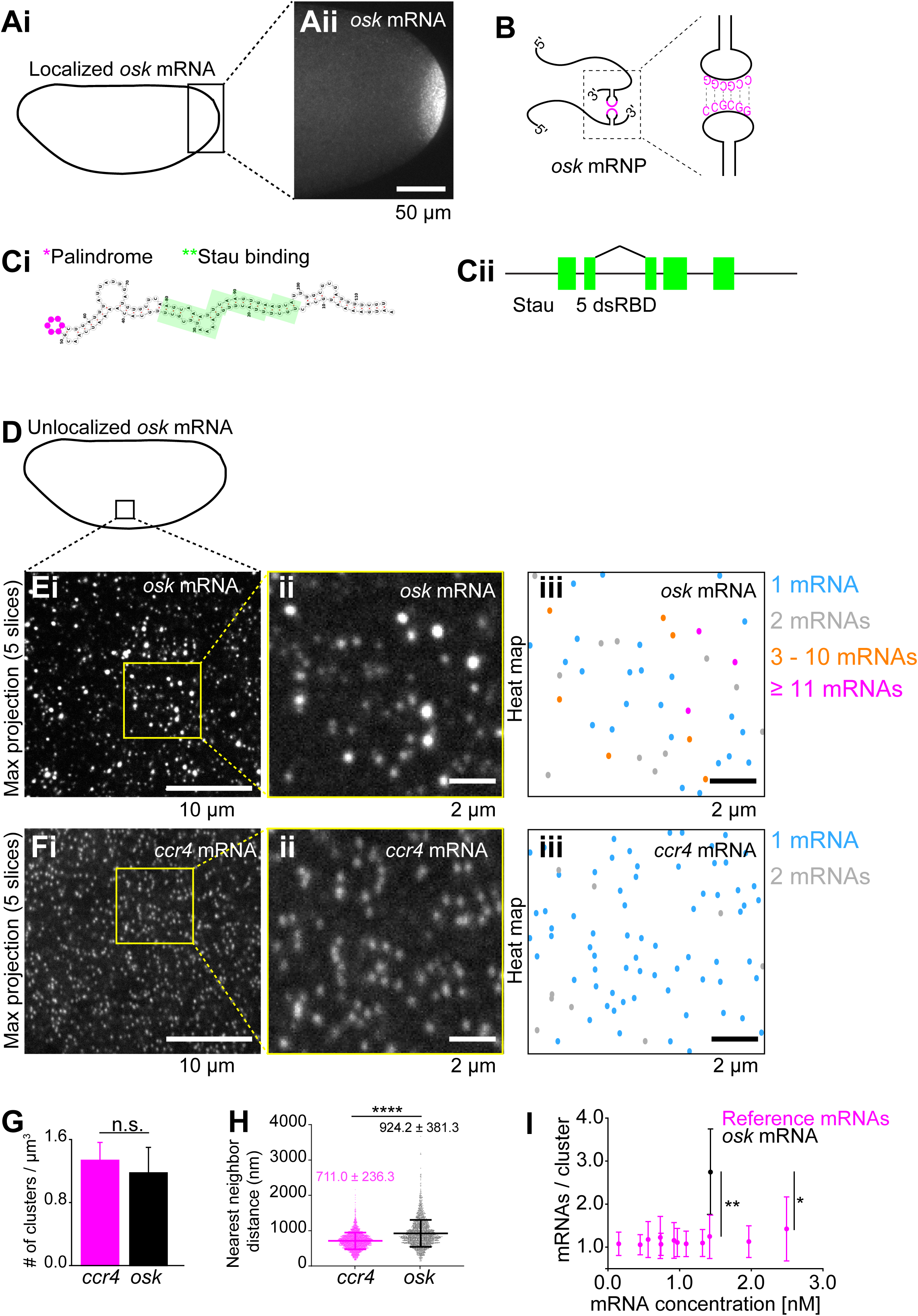
Applying smFISH to examine *osk* mRNA clustering in early embryos. **Ai,ii** Schematic of an embryo depicting the location of localized *osk* mRNA (i) accumulated at the posterior pole stained with smFISH (ii). **B** Schematic illustrating dimerization of *osk* mediated by an RNA palindrome. The palindrome (CCGCGG) is exposed atop of a long RNA stem (magenta) which facilitates intermolecular base pairing of *osk* mRNAs and enables its hitchhiking^15,38^. **Ci,ii.** A diagram showing the position of five dsRBDs (green boxes) of the *Drosophila* Stau^78^ (i) and the position Stau binding to the *osk* stemloop structure (green boxes). The nucleotides of the *osk* palindrome are labeled in magenta^38^ (ii). The stemloop structure comprised nucleotides 714–827, which is the conserved region within the dimerization-competent portion of the *osk* 3′UTR defined previously^38^. **D** Schematic of an embryo depicting an approximate location of a 3D region of interest (ROI) used to analyze clustering of unlocalized *osk* mRNA in Ei-iii. **Ei-iii.** Maximal projection of five consecutive optical sections of an ROI taken from the somatic region of the embryo (Ai), spaced 150 nm apart (i). Yellow ROI: zoomed in image shown in (ii) with an accompanying heat map image showing the distribution of *osk* mRNA clusters of different sizes (iii). **Fi-iii** Maximal projection of five consecutive optical sections spaced 150 nm apart take on an embryo stained with smFISH probes hybridizing to *ccr4* mRNA (i). Yellow ROI: zoomed in image (ii) with an accompanying heat map image (iii). **G** Number of fluorescent *ccr4* or *osk* clusters per µm^3^ volume (density). Mean±STDEV of five and seven 3D ROIs is plotted. Statistical significance: two-tailed *t*-test. n.s.: not significant. **H** Shortest nm distance between neighboring *ccr4* or *osk* clusters. Mean±STDEV of 2999 and 2074 *cc4* and *osk* clusters, respectively were analyzed. Statistical significance: two-tailed *t*-test. ****: P < 0.001. **I** Average size of *osk* mRNA clusters (black circle and error bar) and of 12 reference mRNAs (*aret, ccr4*, *CG18446*, *eIF4G2*, *gcl*, *orb, Pi3K*, *p53*, *pum*, *shutdown*, *sra* and *tao1;* magenta circles and errors bars) plotted against their respective somatic concentrations. The data for the reference mRNAs were determined previously^47^. All differences in cluster sizes between *osk* (black circles) and reference mRNAs (magenta circles) are statistically significant, with the P values ranging from 0.01 (*) to 0.004 (**; two-tailed *t*-test).

In addition to RNA sequence features, RNA concentration and RBPs can also enhance mRNA clustering driven by CSs. For instance, dimerization of the MLV RNA is thought to occur during transcription, where local viral RNA concentration is high^39^. Likewise, HIV-1 RNA dimerization frequently takes place upon RNA localization to the plasma membrane and requires the viral Gag protein^40,41^, while clustering of *BNI1*, *SPA2*, *osk* and *bcd* mRNA similarly depends on their respective RBPs^22,23,42–44^. Together, these observations suggest that RNA concentration and RBPs are important for establishing efficient CS-dependent intermolecular base pairing to promote RNA clustering. However, how these factors enhance mRNA clustering in vivo has not been systematically investigated.

Here, we dissect the contribution of mRNA concentration, RBPs, and intermolecular base pairing to the clustering of the *Drosophila osk* mRNA. *osk*’s ability to engage in intermolecular base pairing has been experimentally validated both in vitro and in vivo. Specifically, *osk* contains an exposed palindrome (CCGCGG) positioned at the apex of a stem within its 3′ untranslated region (UTR), which mediates intermolecular base pairing between *osk* mRNAs^15,38^ (Fig. 1B). This palindrome enables one *osk* mRNA to “hitchhike” on another during its transport to the posterior pole^15,38^ (Fig. 1B). Mutations that disrupt the palindrome abolish dimerization of *osk* 3′UTR in vitro and posterior localization of the mRNA in vivo^15^. Furthermore, *trans* compensatory mutations revealed that dimerization of *osk* occurs through direct intermolecular base pairing in vivo^15^, highlighting the palindrome as a critical *cis*-acting element required for *osk* mRNA localization. At the posterior pole, hundreds of *osk* mRNA copies oligomerize within founder granules^3,45–47^ wherein *osk* palindrome remains engaged in intermolecular based pairing^47^, acting as a “interaction sticker”^43^ that supports *osk* oligomerization in vivo.

While distinct RBPs promote *osk* mRNA clustering^42–44^, in this study we examined the contribution of the double stranded RBP (dsRBP) Staufen (Stau) as a model to evaluate the contribution of RBPs to mRNA clustering. We focused on Stau for several key reasons. First, Stau is essential for *osk* mRNA localization to the posterior pole and in its absence, *osk* fails to localize^48–50^. Since intermolecular base pairing between *osk* mRNA also enhances *osk* localization to the posterior^15^, we reasoned that Stau may contribute to dimerization of *osk* as well. Second, Stau is a dsRBP^51^ that specifically recognizes the stem structure that carries the palindrome within the *osk* 3′UTR (Fig. 1Ci, green segments)^48,52^. Mutations that disrupt base pairing within this stem abolish Stau binding, while compensatory mutations that restore base pairing also restore Stau binding^48^. Moreover, mutations of the first two cytosines in the palindrome disrupt *osk* dimerization in vitro and its localization in vivo^15^, but preserve Stau binding^43^. Therefore, Stau association with the stem that carries the palindrome does not require prior dimerization. This may allow Stau to directly promote initial RNA dimerization steps. Lastly, *Drosophila* Stau contains five RNA-binding domains (Fig. 1Cii)^53^, suggesting that it may function as a molecular bridge, linking RNA monomers and promoting their assembly as well as stabilizing intermolecular base pairing interactions required for oligomerization^22^.

Our experiments revealed that high mRNA concentration, double-stranded RNA-binding protein (dsRBP) Stau, and intermolecular base pairing mediated by the RNA palindrome synergize to optimize *osk* clustering, with oligomerization being more sensitive to perturbations in these factors than dimerization. Stau and the palindrome lower the mRNA concentration threshold required for dimerization, while the palindrome promotes heterotypic interactions between endogenous *osk*and a reporter mRNA bearing the *osk* 3′UTR, providing an important determinant to compositional specificity for mRNA clustering.

Finally, computational analyses identify additional early embryonic mRNAs containing similar palindromes to the one found in *osk*, suggesting that intermolecular base pairing may represent a broader principle of mRNA organization. Given that more than 2,300 mRNAs localize within the early embryo^54^, intermolecular base pairing driven by exposed RNA palindromes could represent a more widespread mechanism of mRNA organization during early *Drosophila* development.

## Results

### Applying smFISH to examine *osk* mRNA clustering in early embryos

We examined the effect of *osk* mRNA concentration, dsRBP Stau and the RNA palindrome on clustering of unlocalized *osk* in early embryos. To this end we combined structured illumination microscopy, a super-resolution approach, with single molecule fluorescent *in situ* hybridization (smFISH) in which we employed 48 smFISH probes to label the *osk* sequence (Fig. 1D, Ei,ii, Suppl. Table 1, Materials and Methods). This allowed us to detect individual and clustered *osk* mRNAs, which would be more difficult if we examined localized *osk* within founder granules at the posterior pole (Fig. 1Ai,ii), where enhanced mRNA clustering occurs^46^.

We quantified the absolute number of *osk* mRNAs within each diffraction-limited smFISH spot, which we termed *osk* cluster size, and *osk* mRNA concentration for each embryo, as we have done previously^4,47,55^ (Materials and Methods). Fitting the distribution of fluorescent puncta to a double-Gaussian curve (Suppl. Fig. 1A, black circles and line) revealed an *osk* population with two mean fluorescence intensities: X_0_ = 154.3±2.9 arbitrary unites (a.u.), and a second mean intensity that was twice the value of X_0_ (X_1_ = 277.2±25.0; P<0.0001, R^2^:0.96; Suppl. Table 1, Materials and Methods).

Importantly, the intensity of the first peak closely matched the value of 154.9±4.6 a.u. determined in embryos in which *osk* mRNA levels were reduced *via* RNAi, whose expression was driven by a GAL4/UAS system under a strong maternal *alpha-tubulin* (*matα*) promoter^56^ (referred to as “RNAi/*matα*”; P<0.0001, R^2^:0.97; Suppl. Table 1). We have confirmed that the reduction of *osk* expression in these embryos was strong (Suppl. Fig. 1B).

Moreover, this value also closely matched the mean fluorescence intensity of 153.0±4.6 a.u. recoded in embryos lacking Stau expression (referred to as “*stau*^*1*^*/stau*^*2*^”; P<0.0001, R^2^:0.95; Suppl. Table 1). We confirmed that the *stau*^*1*^*/stau*^*2*^ embryos failed to localize *osk* at the posterior pole (Suppl. Fig. 1C) and did not hatch (Suppl. Fig. 1D), consistent with previous observations^49^. Also, the fact that the average intensity of the second Gaussian was twice the intensity of the first population is consistent with the second population being dimers of the first (Suppl. Table 1). Therefore, our approach robustly and reproducibly detected single *osk* mRNAs across different genetic backgrounds *in vivo*, and moreover, enabled us to quantify the discrete number of *osk* mRNAs in oligomerized clusters.

To assess the contribution of the *osk* palindrome to mRNA clustering in vivo, we analyzed the previously characterized engineered mRNAs in which the coding sequence (CDS) of eGFP was fused to the *osk* 3′UTR carrying either the WT *osk* CCGCGG palindrome (termed *eGFP-WT*) or mutated AAGCGG sequence (termed *eGFP-AA*; see also Fig. 2Diii,Eiii)^15^. We used 32 smFISH probes targeting the eGFP sequence to detect these transgenes and differentiate them from endogenous *osk* mRNA (Suppl. Table 1). Fitting the distribution of *eGFP-WT* and *eGFP-AA* fluorescent puncta to a Gaussian model revealed a population corresponding to a single mRNA for each transgene with nearly identical mean fluorescence intensities of 51.7±1.4 and 50.0±1.6 a.u., respectively (P<0.0001, R^2^:0.95 and P<0.0001, R^2^:0.97 for *eGFP-WT* and *eGFP-AA*, respectively; Suppl. Table 1), highlighting the robustness and reproducibility of single mRNA detection by smFISH in vivo.

**FIGURE 2:**
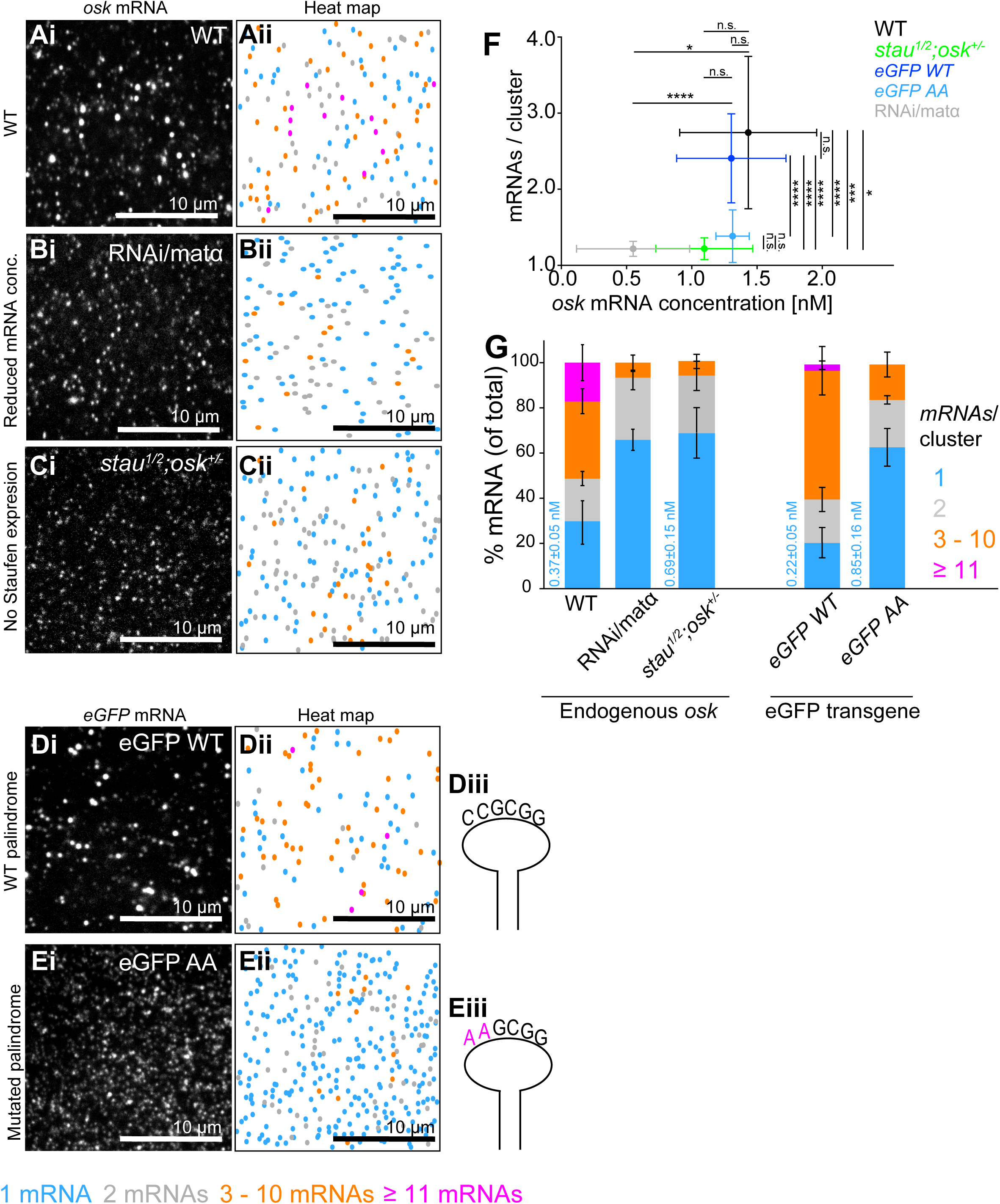
Synergistic action of high *osk* mRNA concentration, Stau expression and the RNA palindrome drive *osk* clustering. **A-Ei-iii.** Maximal projection of five consecutive optical sections spaced 150 nm apart of a WT (Ai), RNAi/*matα* (Bi), *stau*^*1*^*/stau*^*2*^*;osk^+/-^*(Ci) embryos and those expressing *eGFP-WT* (Di) and *eGFP-AA* (Ei) transgenes. Corresponding heat maps depicting the spatial distribution of *osk* (Aii,Bii,Cii), *eGFP-WT* (Dii) and *eGFP-AA* (Eii) mRNA clusters are shown. Schematic of the WT (Diii) and mutated (Eiii) exposed RNA sequence within the *osk* stemloop. **F** Dependence of the average *osk*, *eGFP-WT* and *eGFP-AA* mRNA cluster size on its concentration (nM). Data represents mean±STDEV from seven WT, four RNAi/*matα*, seven *stau*^*1*^*/stau*^*2*^*;osk^+/-^* embryos, 13 *eGFP-WT* and 12 *eGFP-AA* embryos. Statistical significance: two-tailed *t*-test. *: P = 0.01, ***: P = 0.002 and ****: P < 0.001. n.s.: not significant. **G** Distribution of *osk* mRNA across clusters of different sizes. Bars represent mean±STDEV cluster size from seven WT, four RNAi/*matα*, seven *stau*^*1*^*/stau*^*2*^*;osk^+/-^* embryos, 13 *eGFP-WT* and 12 *eGFP-AA* embryos. Blue numbers: the mean±STDEV molarity of *osk*, *eGFP-WT* and 12 *eGFP-AA* mRNA monomers (n = 7,7, 13 and 12 embryos, respectively) for each genotype is shown.

To further validate the accuracy of our single-molecule quantification, we compared *osk* mRNA concentration measured by smFISH with the value extrapolated from a standard curve. This curve was generated by correlating nanomolar (nM) concentrations of six reference mRNAs, namely *germ cell-less* (*gcl*), *tao1*, *ccr4*, *sra*, *pum* and *eIF4G2*, as measured by smFISH, with their respective expression levels determined by RNA-sequencing data obtained from Flybase^57^. These reference mRNAs were selected because we have previously shown that they predominantly appear as single molecules within the early embryo^4,47^. Moreover, their lower expression levels minimized the potential for crowding, which could otherwise lead to undercounting and diminish the accuracy of mRNA concentration extrapolation.

We observed a strong correlation between mRNA expression levels determined by smFISH and RNA-sequencing for the six reference mRNAs (R^2^=0.72; Suppl. Fig. 1E), indicating that smFISH quantified mRNA concentrations across a range of gene expression levels, consistent with our previous findings^4,55^. By fitting the relationship between RPKM values from RNA-seq^57^ and smFISH-derived concentrations with a linear regression, we estimated the somatic concentration of *osk* mRNA to be 1.59 nM (Suppl. Fig. 1E, green circle). This prediction closely matched the 1.43±0.2 nM concentration measured by smFISH (Suppl. Fig. 1E, magenta circle and errors bars). These results indicated that our approach detected single *osk* mRNAs even in the presence of oligomerized species, allowing reliable quantification of *osk* mRNA concentration and its cluster sizes in vivo.

smFISH revealed that WT embryos formed a heterogeneous population of *osk* mRNA clusters, ranging from monomers to assemblies containing more than 11 *osk* transcripts (Fig. 1Ei-iii). In contrast, *ccr4*, our control mRNA, predominantly formed monomers (Fig. 1Fi-iii) even though *ccr4* clusters exhibited similar densities (defined as the number of clusters per volume unit) and slightly decreased cluster-to-cluster distances than *osk* (Fig. 1G,H). In addition, while the fluorescent intensity of *osk* clusters increased with cluster size, their diameters, measured in nanometers using a line scan, did not (Suppl. Fig. 1Fi,ii). Instead, these diameters were comparable to those recorders for *ccr4* clusters (Suppl. Fig. 1G) and for 100 nm TertraSpeck microspheres (Suppl. Fig. 1H), which were below the diffraction limit of light of our structured illumination microscope. Together, these results supported our conclusion that multiple *osk* mRNAs were organized within a confined, diffraction-limited clusters within the early embryos.

Finally, mRNA clustering was specific to *osk*, as other mRNAs, that expressed at similar or higher concentrations than *osk* did not exhibit comparable clustering behavior (Fig. 1I). Therefore, RNA clustering was an intrinsic property of *osk* mRNA, rather than a phenomenon driven by a limited optical resolution of the microscope.

Collectively, our results established robust and consistent detection of single *osk* mRNA using smFISH across distinct genetic backgrounds, providing strong confidence that the observed differences in *osk* cluster sizes reflect *bona fide* changes in cluster organization.

### High *osk* mRNA concentration and Stau expression increase the likelihood of *osk* clustering

Applying smFISH to WT embryos, we recorded an average *osk* cluster size of 2.74±1.00 mRNAs (Fig. 2Ai,ii,F, black circle and error bars), with 70.2% of *osk* mRNA found in clusters that contain more than one *osk* mRNA (Fig. 2G). Therefore, unlocalized *osk* dimerized and oligomerized in WT embryos, consistent with previous observations^3,4^.

Upon reducing the expression of *osk* by 2.6-fold in RNAi/*matα* embryos, *osk* cluster size also reduced by 2.2-fold, a statistically significant difference (Fig. 2Bi,ii,F, gray circle and errors bars). Importantly, comparable *osk* concentrations yielded similar cluster sizes across different genetic backgrounds (Suppl. Fig. 1B, Suppl. Fig. 2A,B), indicating that mRNA concentration rather than genetic background per se drove changes in *osk* clustering in RNAi/*matα* embryos.

In *stau*^*1*^*/stau*^*2*^ embryos, generated by crossing of flies carrying two distinct mutations in the *stau* gene, the average *osk* cluster size similarly decreased from 2.74±1.00 mRNAs in WT early embryos to 1.61±0.12 mRNAs in *stau*^*1*^*/stau*^*2*^ embryos, a statistically significant change (Suppl. Fig. 3A). However, loss of Stau also led to a 64% increase in total *osk* mRNA levels (Suppl. Fig. 3A,B). The mechanism behind this increase is unclear, however it may reflect the Stau-mediated mRNA decay recorded in mammalian cells^58^. Therefore, to evaluate the role of Stau in *osk* clustering without the accompanying effect of *osk* mRNA concentration change, we combined the *stau*^*1*^*/stau*^*2*^ mutations with the *osk* gene deficiency *Df(3R)p^XT^*^*103* 49^ (hereafter “*stau*^*1*^*^/^*^2^*;osk^+/-^*”), which resulted in *osk* mRNA expression levels comparable to those recorded in WT embryos (Fig. 2F, green circle and errors bars, Suppl. Fig. 3A,B). Under these conditions, the average *osk* cluster size decreased by 55.5% from 2.74±1.00 recorded in WT embryos to 1.22±0.14 in *stau*^*1*^*^/^*^2^*;osk^+/-^*embryos, a statistically significant change (Fig. 2Ci,ii, F). The distribution of *osk* cluster sizes paralleled that observed in RNAi/*matα* embryos (Fig. 2G) despite *osk* concentration being approximately two-fold higher than the one recorded for RNAi/*matα* embryos (Fig. 2F, green versus gray circle). These data indicated that high mRNA concentration and Stau expression both augmented *osk* clustering.

Next, we examined how *osk* cluster sizes depended on RNA concentration and Stau expression. Focusing first on oligomers, we observed that the fraction of oligomerized *osk* sharply decreased by 7.8- to 8.7-fold from 51.3 % in WT to 6.6% and 5.9±5.4% in RNAi/*matα* and *stau*^*1*^*^/^*^2^*;osk^+/-^*embryos, respectively (Fig. 2G). Similar dependence of cluster size distribution on *osk* concentration was recorded in embryos with different genetic backgrounds (Suppl. Fig. 2B). Commercially available overexpression lines generated only a modest 2.7-fold upregulation of *stau* mRNA levels (Suppl. Fig. 3E), limiting our ability to further examine whether changes in Stau protein levels augment *osk* clustering. Instead we investigated *stau*^*1*^*/stau*^*2*^ embryos in which the proportion of dimerized *osk* doubled, from 18.9±3.8% in WT early embryos to 40.7±1.2% in *stau*^*1*^*/stau*^*2*^ embryos (Suppl. Fig. 3Ci,ii,D). This result revealed that the transition of dimers into oligomers was impaired in the absence of Stau. Therefore, high RNA concentration and Stau both promoted *osk* mRNA oligomerization.

We then examined *osk* monomers and dimers. Reduced mRNA levels doubled the fraction of *osk* monomers in RNAi/*matα* embryos (Fig. 2Bi,ii, G) consistent with a decreased probability of intermolecular encounters and *osk* dimerization at lower transcript abundance.

However, in *stau*^*1*^*^/^*^2^*;osk^+/-^* embryos *osk* mRNA expression level was comparable to the one recorded in WT embryos (Fig. 2F, green and black circles, Suppl. Fig. 3A,B), allowing us to compare changes in *osk* monomers and dimers directly. Specifically, we observed that the concentration of *osk* monomers doubled from 0.37±0.05 nM to 0.69±0.15 nM recorded for WT and *stau*^*1*^*^/^*^2^*;osk^+/-^*embryos, respectively (Fig. 2G, blue numbers). A similar increase in monomer concentration was recorded in *stau*^*1*^*/stau*^*2*^ embryos, which exhibited increased *osk* expression relative to WT (Suppl. Fig. 3A-D). Therefore, dsRBP Stau promoted *osk* clustering by lowering the concentration threshold required for its dimerization in vivo.

Taken together, our analysis revealed that *osk* dimerization is governed by both *osk* mRNA concentration and dsRBP Stau and that the formation of higher-order oligomers is more acutely sensitive to perturbations in these factors. This differential sensitivity suggests that dimerization and oligomerization rely on distinct combinations of intermolecular interactions. Notably, Stau expression approximately doubles the probability of *osk* dimerization at lower expression levels, indicating that it lowers the concentration threshold required for mRNA dimerization.

### *osk* palindrome enhances mRNA dimerization and oligomerization

Next, we examined the contribution of *osk* palindrome to mRNA clustering. In vitro, dimerization of the RNA derived from *osk* 3′UTR depends on a palindrome presented within a stemloop structure (Fig. 1B)^15,38^ and mutations that disrupt this sequence also abolish *osk* localization to the posterior pole in vivo^15^. Additionally, substitution of the first two cytosines of the palindrome to adenosines increases the apparent dissociation constant (*K_d_*) for *osk* 3′UTR dimerization by threefold, from 90 nM observed for WT *osk* 3′UTR carrying the CCGCGG palindrome to ∼270 nM observed for *osk* 3′UTR carrying the mutated AAGCGG variant^15^.

To asses this role in vivo, we analyzed clustering of *eGFP-WT*, carrying the CCGCGG palindrome and *eGFP-AA* carrying the mutated AAGCGG sequence^15^ (Fig. 2Di-iii, Ei-iii). However, *osk* 3′UTR performs an essential noncoding function during oogenesis and females that do not express *osk* do not lay eggs^59,60^. Although the UAS-Gal4/*nanos* promoter system is active throughout oogenesis^61^, it did not drive sufficient expression of *eGFP-WT* and *eGFP-AA* transgenes in *osk^A^*^87^*/Df(3R)p^XT^*^*103*^ females, which lacked endogenous *osk* RNA expression, to restore egg laying (Suppl. Fig. 4A,B).

Instead, we expressed both eGFP transgenes using the UAS-Gal4/*matα* system^56^ and detected them alongside endogenous *osk* using spectrally distinct smFISH probes targeting their respective CDSs. Importantly, both *eGFP-WT* and *eGFP-AA* exhibited similar expression levels (Suppl. Fig. 4C,D), allowing us to compare the dimerization and oligomerization potential of *eGFP-AA* against *eGFP-WT*. We observed that while *eGFP-WT* mRNA localized at the posterior pole (fold enrichment of 1.96±0.36, defined as the ratio of eGFP fluorescent levels between the posterior and outside, Suppl. Fig. 4Ei,ii, G,H), *eGFP-AA* mRNA displayed minimal localization (fold enrichment of 1.20±0.15, Suppl. Fig. 4Fi,ii, G,H), indicating that the intact palindrome is necessary for efficient localization of *osk* to the posterior, in agreement with published observations^15^.

Clustering of *eGFP-WT* mirrored that of endogenous *osk* (Fig. 2Ai,ii,Di,ii,G): 79.4% of unlocalized *eGFP-WT* was found in clusters that contained more than one *eGFP-WT* mRNA (Fig. 2G), yielding an average cluster size of 2.40±0.58 mRNAs (Fig. 2F, dark blue circle and error bars). This trend remained even after *eGFP-WT* clusters that associated with the endogenous *osk* mRNA were removed from the analysis (Suppl. Fig. 4I, also see Fig. 3). Therefore, like endogenous *osk*, unlocalized *eGFP-WT* mRNA efficiently dimerized and oligomerized.

**FIGURE 3:**
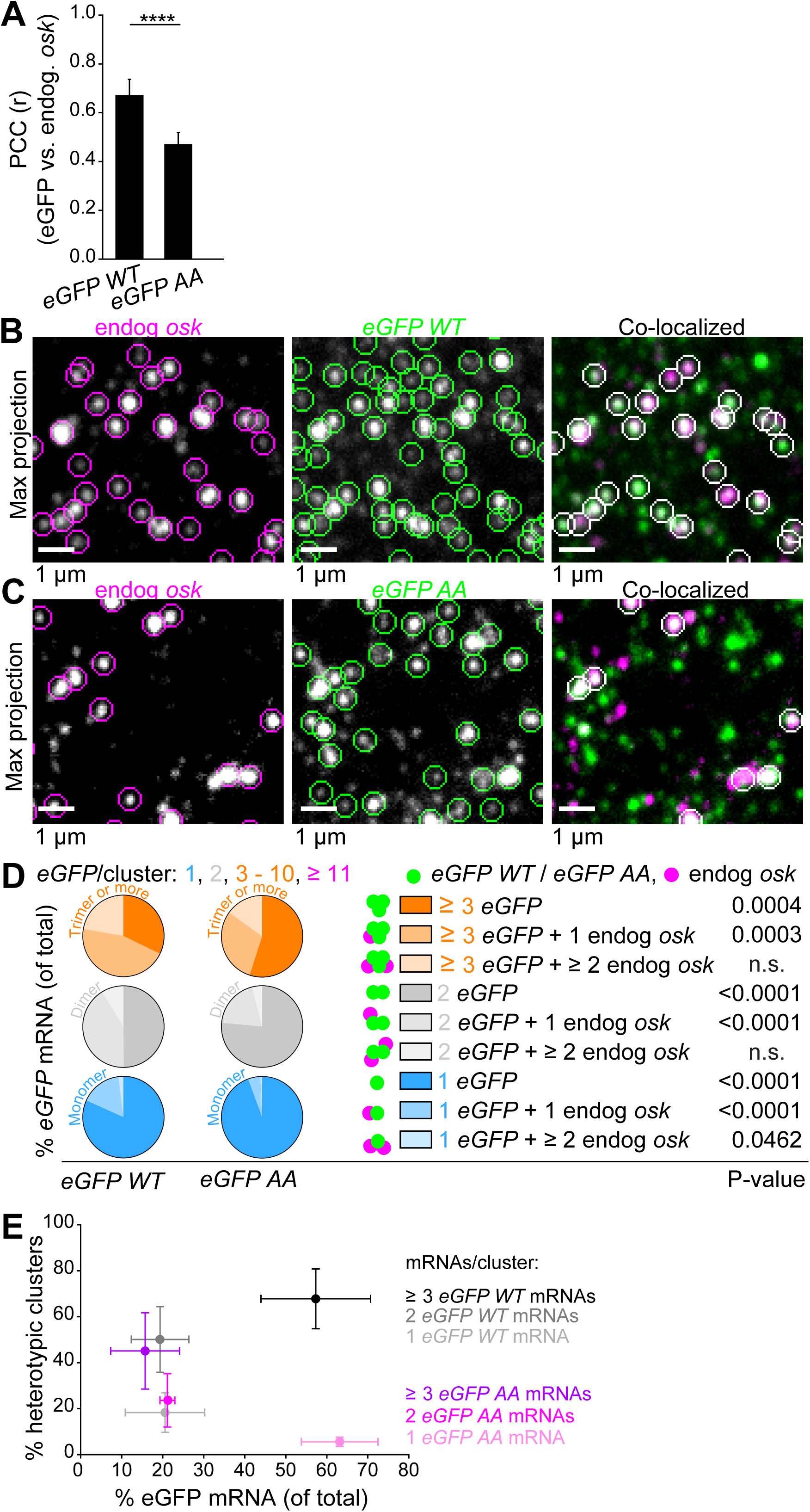
*osk* palindrome is a major determinant of heterotypic *osk* mRNA clustering. **A** Co-localization between endogenous *osk* and *eGFP-WT* or *eGFP-AA,* quantified with Pearson’s correlation coefficient (PCC). Mean±STDEV of PCC values from 13 *eGFP-WT* and 12 *eGFP-AA* embryos are shown. Statistical significance: two-tailed *t*-test. ****: P < 0.001. **B,C** Representative images showing co-localization between endogenous *osk* (green) and *eGFP-WT* (C) or *eGFP-AA* (D) (magenta) with a 250 nm cutoff. Maximal projection of 5 consecutive optical sections of an ROI, spaced 150 nm apart are shown. **D** Pie charts show the composition of each *eGFP* cluster category based on its association with endogenous *osk* mRNA. For example, 20.6±9.7% of *eGFP-WT* spots display the intensity expected for a single *eGFP-WT* mRNA (“monomers”; dark blue in the bar chart). Co-localization analysis using spot detection revealed that 90.4±5.4% of these were true monomers (dark blue in the pie chart and legend), whereas 8.7±4.4% and 1.0±1.2% co-localized with endogenous *osk* dimers and oligomers, respectively (lighter blue shades in the pie chart and legend). Differences in heterotypic cluster size between *eGFP-WT* and *eGFP-AA* were evaluated using a two-tailed *t*-test. **E** Fraction of *eGFP-WT* and *eGFP-AA* clusters with heterotypic composition per given *eGFP-WT* (black) and *eGFP-AA* (magenta) cluster size. For example, of the 63.1±9.3% of *eGFP-AA* mRNA (magenta) identified as monomeric, 5.6±2.0% were heterotically associating with endogenous *osk*, while 20.6±9.7% of *eGFP-WT* mRNA identified as monomeric (black), 18.3±8.6% were heterotipically associating with endogenous *osk*. Mean±STDEV from 13 *eGFP-WT* and 12 *eGFP-AA* embryos is shown.

In contrast, *eGFP-AA* showed impaired clustering which mirrored that of *osk* in RNAi/*matα* embryos where the *osk* levels were reduced (Fig. 2Bi,ii,E-G). Only 36.9% of unlocalized *eGFP-AA* was found in clusters containing more than one mRNA (Fig. 2G), yielding an average size of 1.31±0.12 mRNAs, a statistically significant decrease compared to *eGFP-WT* (Fig. 2F, light blue circle and error bars). Notably, this reduction occurred despite similar expression of *eGFP-AA* and *eGFP-WT* (1.4±0.3 nM and 1.3±0.4 nM, respectively, Fig. 2F, Suppl. Fig. 4C,D). Consequently, the concentration of single *eGFP-AA* molecules nearly quadrupled, from an 0.22±0.05 nM for *eGFP-WT* mRNA to 0.85±0.16 nM for *eGFP-AA* mRNA (Fig. 2G, blue numbers). This trend remained after *eGFP-AA* clusters that associated with the endogenous *osk* mRNA were removed from the analysis (Suppl. Fig. 4I, also see Fig. 3). Notably, the magnitude of this decrease was comparable to the ∼threefold reduction in dimerization potential for the *osk* 3′UTR carrying the mutated AAGCGG palindrome relative to the WT palindrome recorded in vitro^15^.

These results revealed that *eGFP-AA* mRNA with a disrupted palindrome still dimerized, albeit inefficiently. In contrast, the intact palindrome of *eGFP-WT* greatly enhanced dimerization, effectively quadrupling its probability. Thus, the *osk* palindrome lowered the concentration threshold required for dimerization and subsequent oligomerization.

### The RNA palindrome is a major determinant of heterotypic *osk* mRNA clustering

To further assess the role of the palindrome in clustering of *osk*, we analyzed the ability of *eGFP-WT* and *eGFP-AA* transgenes to heterotypically dimerize and oligomerize with the endogenous *osk* using spectrally distinct smFISH probes targeting their respective CDSs.

Applying Pearson’s Correlation Coefficient (PCC) analysis to evaluate co-localization, we found that *eGFP-WT* better co-localized with endogenous *osk* than *eGFP-AA* (Fig. 3A), indicating that the palindrome enhances the likelihood for heterotypic mRNA clustering.

Next, we analyzed the composition of *eGFP-WT* and *eGFP-AA* mRNA clusters. We quantified the number of *eGFP* monomers, dimers, and oligomers that co-localized with one, two, or more endogenous *osk* mRNAs. A 250-nanometer (nm) cutoff between *eGFP* transgene and endogenous *osk* was used to define co-localization (Fig. 3B,C, Materials and Methods; Suppl. Fig. 5A,B). This inclusive cutoff was based on our previous measurement showing that, in founder granules in embryos, *osk* forms clusters with a diameter of ∼ 100 nm^47^, which corresponds to a spacing between neighboring *osk* clusters of approximately 100–150 nm.

We observed that of the 20.6 ± 9.7% of all *eGFP-WT* monomers (Fig. 2G), 81.7 ± 8.6% were true monomers, with the remaining fraction co-localizing with one or more endogenous *osk* mRNAs (Fig. 3D, blue pie chart). Similarly, of the 19.3 ± 7.0% and 60.1% of all *eGFP-WT* dimers and oligomers, respectively (Fig. 2G), 49.9 ± 14.3% and 32.2 ± 13.0% formed homotypic clusters, respectively, while the remainder co-localized with endogenous *osk* mRNAs (Fig. 3D, grey and orange pie chart, respectively).

In contrast, of the 63.1 ± 9.3% of all *eGFP-AA* monomers (Fig. 2G), 94.4 ± 2.0% were true monomers, with the rest co-localizing with endogenous *osk* (Fig. 3D, blue pie chart). Moreover, of the 21.1 ± 1.8% and 15.7% of all *eGFP-AA* dimers and oligomers, respectively (Fig. 2G), 76.4 ± 11.6% and 54.9 ± 16.6% dimers and oligomers formed homotypic clusters, respectively, showing statistically significant reduction in heterotypic interaction compared to *eGFP-WT* (Fig. 3D, grey and orange pie chart, respectively). Importantly, this trend persisted even when the co-localization threshold was reduced to 100 nm (Suppl. Fig. 5C,D), indicating that the observed differences were robust to spatial resolution. Therefore, *osk* palindrome promoted heterotypic association between endogenous *osk* and the eGFP chimera.

Interestingly, the probability for heterotypic association applied differently to monomers, dimers and oligomers. Specifically, even though monomers represented the majority (63.1±9.3%) of *eGFP-AA* mRNAs, fewer than 6% formed heterotypic clusters with endogenous *osk* (Fig. 3E). In contrast, while monomers represented only 20.6±9.7% of *eGFP-WT* transcripts, they were 3.3-fold more likely than *eGFP-AA* mRNAs to form heterotypic clusters with endogenous *osk* (Fig. 3E). Therefore, the palindrome was a potent enhancer of heterotypic dimerization.

In contrast, the fraction of *eGFP-AA* mRNAs that formed heterotypic clusters with endogenous *osk* increased by more than four-fold to 23.5% and 45.1%, respectively, even though these clusters contained less than 40% of all *eGFP-AA* mRNA (Fig. 3E). These findings suggested that heterotypic association of a single *eGFP-AA* with endogenous *osk* is limited but becomes substantially more likely once *eGFP-AA* dimers and oligomers have formed. We observed a similar, although more pronounced trend with the *eGFP-WT* mRNA (Fig. 3E).

These data suggested that the ability of *osk* monomers to form heterotypic interactions were dominated by the *osk* palindrome, whereas heterotypic interactions involving dimers and oligomers were enhanced by additional interactions that may be driven by RBPs as well as other intermolecular base pairing interactions. This model is consistent with our observation that oligomerization of endogenous *osk* is mediated by multiple types of interactions, involving both RNAs and proteins (Fig. 2).

Together, our results demonstrated that the RNA palindrome not only enhanced dimerization and oligomerization of *osk* mRNA but also promoted its heterotypic organization with mRNAs derived from distinct genetic loci.

### RNA palindromes may spatially organize other mRNAs in the early *Drosophila* embryo

Finally, we asked whether other *Drosophila* mRNAs may contain palindromes similar to *osk*. To address this, we identified all palindromes in transcripts of early embryos (NC 1 to 14) using an in-house script we described previously^25^. Following the *osk* example, we looked for GC-rich palindromes with the minimal nucleotide length of six nucleotides. We analyzed 11657 non—localizing transcripts (termed “ubiquitous”) representing isoforms of 4304 genes (Suppl Table 2). Because *osk* localizes to the posterior, we reasoned that a similar palindrome-driven dimerization mechanism may contribute to the localization of other posterior enriched mRNAs. Therefore, we separately analyzed 211 transcripts representing isoforms of 59 mRNAs previously identified as germ granule-enriched (termed “posterior localized”)^62^ (Suppl Table 2).

We observed that the total number of palindromes per transcript scaled linearly with transcript length (Suppl. Fig. 6A) and that palindromes were modestly abundant across transcripts regardless of posterior localization (Suppl. Fig. 6B). Therefore, the presence of the palindrome was not predictive of an mRNA’s ability to enrich at the posterior pole. GC-rich palindromes, defined here as having a GC content equal or greater than 50%, accounted for less than half of the total palindrome density, indicating that palindromes were more frequently AT-rich than GC-rich (Suppl. Fig. 6B).

Because the *osk* palindrome is both GC-rich and presented atop an RNA stem-loop (Fig. 1B)^15,38^, we searched for GC-rich palindromes that resided within such RNA structures (“Materials and Methods”). As the full secondary structure of the *Drosophila* transcriptome was not experimentally available, we approximated RNA structure using Vienna RNAfold. Due to the symmetrical nature of palindromes, a palindrome whose first half is embedded but second half is exposed is not able to engage in intermolecular dimerization. This is because the base pairing partner of the second half is the first half. Therefore, to gain a more accurate understanding of how many palindromes may be able to dimerize, we considered the maximum number of consecutive nucleotides that were exposed in both palindrome halves termed the “number of base-pairable positions” or “bp score” (Fig. 4 Ai; Suppl. Table 2).

**FIGURE 4:**
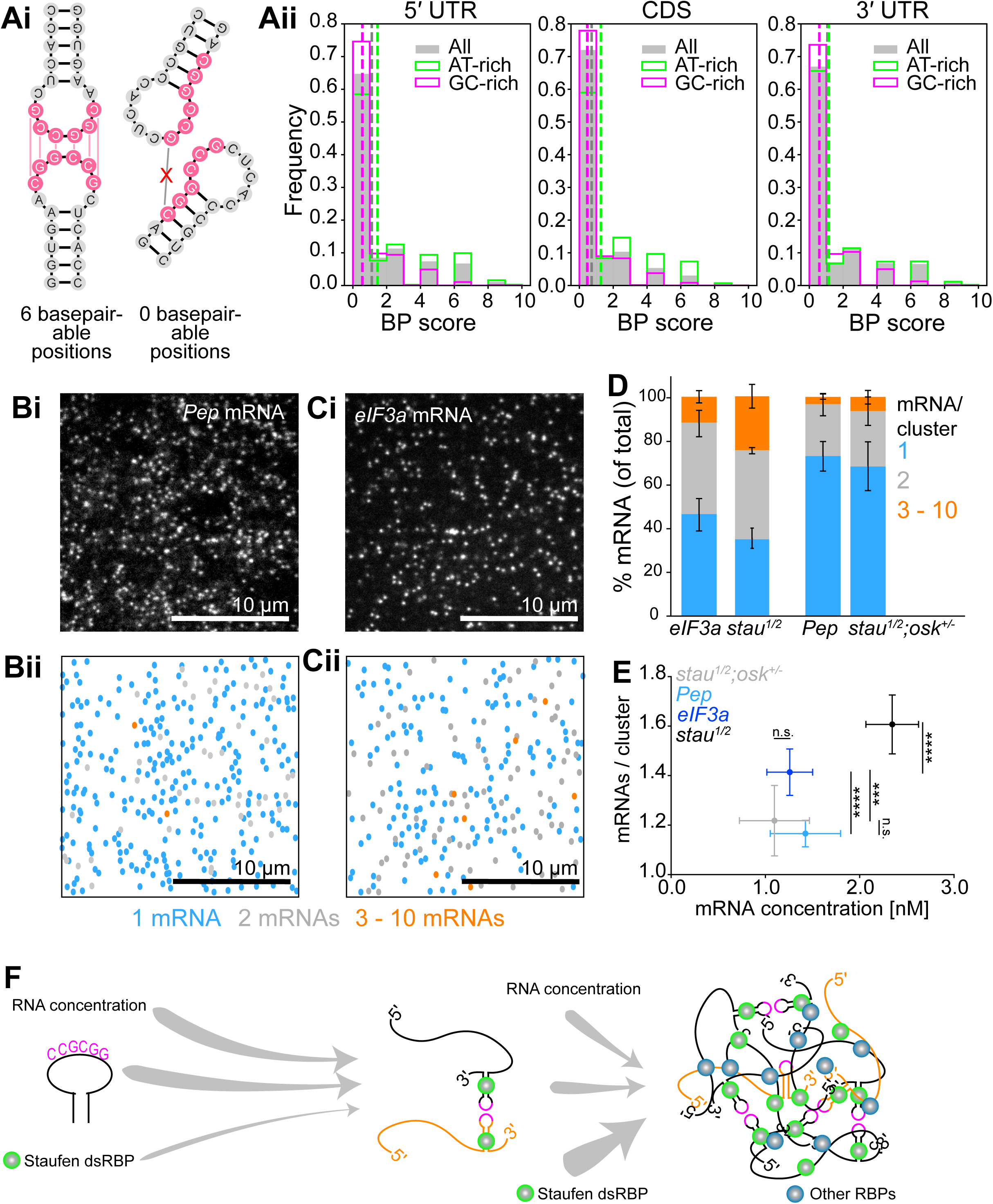
RNA palindromes may spatially organize other mRNAs in the early *Drosophila* embryo. **Ai,ii** Schematic of two palindromes presented atop an RNA stem-loop with 6 or 0 base-pairable positions (Materials and Methods) (i). Histogram of the distribution of all (gray), AT-rich (green) and GC-rich (magenta) palindromes with a given number of base-pairable positions identified within the 5′UTR, CDS and 3′UTR of ubiquitous and posterior localized transcripts expressed in the early *Drosophila* embryo (ii). **Bi-Cii** Maximal projection of five consecutive optical sections of an ROI, spaced 150 nm apart of *Pep* mRNA (Bi) and *eIF3a* mRNA (Ci) with corresponding heat maps depicting the spatial distribution of *Pep* and *eIF3a* mRNA clusters (Bii, Cii). **D** Distribution of *Pep* and *eIF3a* mRNA across clusters of varying sizes. For each cluster category, mean±STDEV of 14 embryos per mRNA is shown. Data for *stau*^*1*^*/stau*^*2*^ and *stau*^*1*^*/stau*^*2*^*;osk^+/-^* are from Suppl. Fig. 3D and Fig. 2G, respectively. **E** Dependence of average *osk* mRNA cluster size of on its concentration (nM). For *Pep* and *eIF3a* mRNAs, data represent mean±STDEV from 14 embryos. Data for *stau*^*1*^*/stau*^*2*^ and *stau*^*1*^*/stau*^*2*^*;osk^+/-^* are from Suppl. Fig. 2A and Fig. 2F, respectively. Statistical significance: two-tailed *t*-test. ***: P = 0.001. ****: P < 0.001. n.s.: not significant. **F** A model showing how the RNA palindrome, elevated mRNA concentration, and Stau dsRBP synergize to promote *osk* dimerization and subsequent oligomerization. While Stau doubles the likelihood of *osk* dimerization (thinner arrow), the palindrome quadruples it (thicker arrow) indicating that Stau and the palindrome lower the critical concentration of *osk* mRNA required for dimerization. Moreover, the palindrome also markedly increases the likelihood of heterotypic *osk* association (depicted as interacting black and orange RNAs), establishing it as the major contributor to heterotypic *osk* mRNA organization. Dimerization and oligomerization may be further supported by additional RBPs (blue circles) and intermolecular base pairing interactions not studied here.

We observed that palindromes with at least six base-pairable positions were mostly AT-rich, while those resembling the GC-rich palindrome of *osk* were extremely rare (Fig. 4Aii). Looking at both ubiquitous and posterior localized mRNAs and their isoforms, we identified 60 (1.4%) mRNAs representing 155 (1.3%) isoforms that contained exposed, GC-rich palindromes (Suppl. Table 3). The majority of these palindromes were found in the CDS (in 38 out of 60 mRNAs and 101 out of 155 isoforms), while the 3′UTR had the least (in 9 out of 60 mRNAs and 19 out of 155 isoforms; 4Aii; Suppl. Table 3). While none of these mRNAs were simultaneously highly expressed and predicted Stau targets (Suppl. Table 3), two mRNAs with GC-rich palindromes found in the CDS, *Pep* and *eIF3a*, nevertheless stood out as their expression levels recorded by RNA-seq were the most similar to the one recorded for *osk* (Suppl. Table 3).

smFISH analysis revealed that neither *Pep* nor *eIF3a* exhibited the strong clustering behavior characteristic of *osk*. Even so, 42.5±6.8 % and 11.3±6.8 % of *eIF3a* dimerized and oligomerized, respectively (Fig. 4B-D). The distribution of *eIF3a* cluster sizes was comparable to *osk* in *stau*^*1*^*^/^*^2^ embryos (Fig. 4D), which was noteworthy given that *eIF3a* expression was approximately two-fold lower than that of *osk* in *stau*^*1*^*^/^*^2^ embryos (Fig. 4E). In contrast, only 24.3±6.0 % and 2.8±1.5 % of *Pep* dimerized and oligomerized, respectively (Fig. 4D), displaying a similar cluster size distribution observed for *osk* in *stau*^*1*^*^/^*^2^*;osk^+/-^* embryos (Fig. 4D,E). These data suggested that *eIF3a* mRNA may be a *bona fide* candidate transcript, whose spatial organization, although moderate, was driven by intermolecular base pairing. Importantly, the early embryo may not present an environment that would support abundant clustering of *eIF3a* as observed for *osk*. For example, *eIF3a* may require distinct dsRBPs, higher mRNA concentration, as well as additional intermolecular base pairing interactions for efficient clustering.

In summary, our transcriptome-wide characterization revealed that while not wide-spread, exposed GC-rich palindromes may not be unique to *osk* mRNA. More than 2300 maternally-deposited mRNAs localize within the early *Drosophila* embryo^54^ where they promote the establishment of early *Drosophila* lineages and embryonic patterning. Exposed GC-rich palindromes may therefore represent an important and previously underappreciated mechanism that contributes to mRNA organization and development of the early *Drosophila* embryo.

## DISCUSSION

### Persistent intermolecular base pairing requires the involvement of high RNA concentration and RBPs

Compared to in vitro studies using the isolated *osk* 3′UTR sequence^15,43^, the conditions that support intermolecular base pairing of *osk* RNA in vivo are more stringent and require the combined action of high RNA concentration, and Stau and the RNA palindrome. This principle closely mirrors RNA dimerization of the HIV-1^63^.

HIV-1 contains a GC-rich palindrome (“GCGCGC” or “GUGCAC” depending on the subtype^64,65^) exposed within the SL1 stem–loop^63^ that promotes HIV-1 RNA dimerization in vitro^66^ and facilitates heterotypic RNA dimerization in cells^67^. However, as with *osk*, the palindrome alone is insufficient for efficient HIV-1 RNA dimerization, and additional factors are required. The HIV-1 nucleocapsid protein 7 (NCp7), which is produced by processing of the larger polyprotein Gag^40,41^ binds to the stem-loop containing the GC-rich palindrome^68,69^, and stabilizes RNA dimers in vitro^69–72^ and in cells^73^. Expression of Gag increases the population of dimeric RNAs by roughly three-fold^73^, paralleling the effect of Stau on *osk* dimerization in embryos (Fig. 2). Thus, NCp7 and Stau appear to act analogously as molecular “glues” that stabilize palindrome-initiated RNA-RNA interactions.

Moreover, in vivo, dimerization of the HIV-1 frequently occurs at the plasma membrane in the presence of the Gag protein^73^ where membrane association likely elevates local RNA concentration. Indeed, many RNA viruses assemble at sites of high RNA density, including transcription sites, the plasma membrane, or spherule invaginations derived from vesicles formed by the mitochondrial outer membrane^39,73,74^. This reinforces our observation that exposed base-pairing sequences and elevated local RNA concentration cooperate with RNA-binding proteins to enable efficient and sustained intermolecular RNA interactions in vivo.

### Multivalency and higher-order mRNA clustering

Despite their similarities, HIV-1 packages a defined RNA dimer into the virion, whereas *osk* transcripts assemble into clusters that may contain many mRNA copies (Fig. 2Ai,ii)^3,46^. This difference likely reflects interactions valency promoted by NCp7 and Stau, other RBPs as well as the RNA itself.

NCp7 contains two zinc finger motifs that primarily bind the stem-loop structure responsible for HIV-1 RNA dimerization, with some binding also detectable with the adjacent stem-loop^40,75–77^. In contrast, Stau contains five dsRBDs (Fig. 1Cii) and binds the palindrome-containing stem with dsRBD 3 (Fig. 3Ci)^48,53,78^. This architecture suggests a model in which Stau stabilizes dimers through dsRBD3-mediated binding, while the remaining four dsRBDs could bridge additional *osk* transcripts, bringing them into proximity and enabling higher-order oligomerization.

Importantly, mutations in the palindrome reduce but do not abolish *osk* dimerization (Fig. 2). This is consistent with in vitro studies showing that disruption of the palindrome does not abolish dimer formation of the isolated *osk* 3′UTR on gels^15,43^. These findings suggest that additional RNA elements may promote intermolecular RNA:RNA interactions even in the absence of *osk* palindrome, in line with recent in vitro RNA phase-separation experiments using the *osk* 3′UTR^43^. Alternatively, the palindrome may mediate initial recognition, whereas other factors, including RBPs, stabilize the interaction. In this scenario, partial disruption of the palindrome would reduce, but not eliminate, dimerization.

Similarly to the mutations of the palindrome, loss of Stau strongly reduced, but did not abolish, *osk* dimerization and oligomerization (Fig. 2Di,ii,Ei,ii), indicating that additional factors contribute to this process. Two RBPs, the polypyrimidine tract-binding (PTB) protein and translational repressor Bruno, contribute to intermolecular *osk* interactions within founder granules at the posterior pole^42–44,79^, and they could compensate for the lost Stau in oligomerization of unlocalized *osk*. Both proteins exhibit multivalent RNA-binding capacity^42,44,79–81^, and Bruno can also dimerize^82^, potentially reinforcing *osk* oligomerization.

Notably, *Drosophila* PTB contains four RNA recognition motifs capable of binding multiple PTB-binding sites within the *osk* 3′UTR^42^. Similarly, Bruno harbors three RBDs, each capable of engaging multiple Bruno response elements *osk*^44,79–81^. Given the multivalent RNA-binding capacity of both PTB and Bruno, together with Bruno’s ability to dimerize^82^ and the presence of multiple binding sites for these proteins within *osk* mRNA, collectively these proteins likely provide an additional interaction layer that reinforces oligomerization of unlocalized *osk*. Thus, *osk* clustering is supported by multivalency at both the RNA and protein levels.

### The palindrome as a determinant of compositional specificity

Interestingly, we observed that *eGFP-AA* mRNAs were 3.3-fold less likely than *eGFP-WT* mRNAs to dimerize with endogenous *osk* (Fig. 3E), despite maintaining the binding sequences for Stau, Bruno or PTB^43^. This indicates that the palindrome itself is a dominant driver of heterotypic mRNA:mRNA recognition. A similar principle operates in *Ashbya gossypii*, where base pairing between *BNI1* and *SPA2* mRNAs establishes compositional specificity of condensates associated with polarized growth^23^. By analogy, our data support a model in which palindrome-mediated base pairing drives initial recognition and dimerization, while RBPs predominantly stabilize and expand assemblies.

Interestingly, a substantial fraction of reporter-derived oligomers remained homotypic (32.2±13.0% for *eGFP-WT* and 54.9±16.6% for *eGFP-AA* (Fig. 3D)), suggesting a possible demixing of oligomers as observed for mRNAs that enrich in *Drosophila* germ granules^4,47^. Alternatively, dimerization and oligomerization may occur early in the *osk* mRNA life cycle. Both PTB and Bruno function in mRNA splicing in the nucleus and shuttle between the nucleus and cytoplasm^80,83,84^. This raises the possibility that *osk* clustering could initiate co-transcriptionally, during or shortly after nascent transcript synthesis. This would mirror transcription-dependent dimerization proposed for the Murine leukemia virus^39^. If clusters form co-transcriptionally at the site of synthesis, such assembly could ensure that endogenous *osk* and reporter alleles remain segregated once transported into the cytosol.

### RNA palindromes as a broader mechanism of mRNA organization

Our computational analysis identified 60 mRNAs expressed in the early embryo that contained palindromes similar to the one found in *osk* (Suppl. Table 3). Subsequent smFISH analysis confirmed that one of these, *eIF3a*, also exhibited light clustering behavior comparable to *osk* in *stau*^*1*^*^/^*^2^ embryos (Fig. 4D). Although none of these candidate mRNAs in were both highly expressed and predicted Stau targets early embryos, these parameters may change during later stages of development. Indeed, several transcripts reach expression levels comparable to or higher than *osk* at later developmental time points (Suppl. Table 3).

Furthermore, efficient clustering of these mRNAs may require distinct double-stranded RNA-binding proteins (dsRBPs) as well as additional intermolecular base-pairing interactions. Notably, the *Drosophila* genome encodes at least 12 dsRBPs^85^, any of which could contribute to the spatial organization of specific transcripts during development. Therefore, the palindrome–mediated mRNA organization may be more widespread in *Drosophila* than previously appreciated. More than 30% of maternally deposited transcripts exhibit spatial organization^54^, where they play critical roles in early developmental patterning^1,86^. Thus, although exposed RNA palindromes are not ubiquitous, they may nonetheless represent an important and previously underappreciated mechanism contributing to mRNA organization in the early *Drosophila* embryo, and potentially in other organisms.

In conclusion, intermolecular base pairing has emerged as a potent mechanism of mRNA organization. We find that although RNA palindromes can robustly promote intermolecular interactions in vivo, the cellular conditions that support such base pairing extend well beyond simple sequence complementarity. Thus, in vivo, robust mRNA clustering require cooperation between multiple RNA motifs and RNA-binding domains. Importantly, such synergy may impose regulatory stringency on intermolecular base pairing in cells, making RNA clustering a controlled and context-dependent process rather than a promiscuous outcome of sequence complementarity, as is often observed in vitro. This increased stringency may be critical to prevent unintended interference with translation, as reported for condensates formed by in vitro–transcribed RNAs^87^.

## ACKNOWLEDGEMENTS

We would like to thank Drs. Sarah Woodson and Andrew Gordus for providing comments on our work. This research was supported in part by grant no. NSF PHY-2309135 to the Kavli Institute for Theoretical Physics (KITP). This research was supported by the NIGMS R35GM142737 grant awarded to TT.

## AUTHOR CONTRIBUTION

TT wrote the manuscript with all authors providing edits. TT wrote the discussion. ZY, provided data and methods sections for Figure 4, AE for Figure 3, ST for Figure 3, and TT for Figure 1 through 4. TT wrote the manuscript with all the authors participated in the revision and editing. TT: conceptualization.

## SUPPLEMENTAL FIGURE LEGENDS

**SUPPLEMENTAL FIGURE 1:**
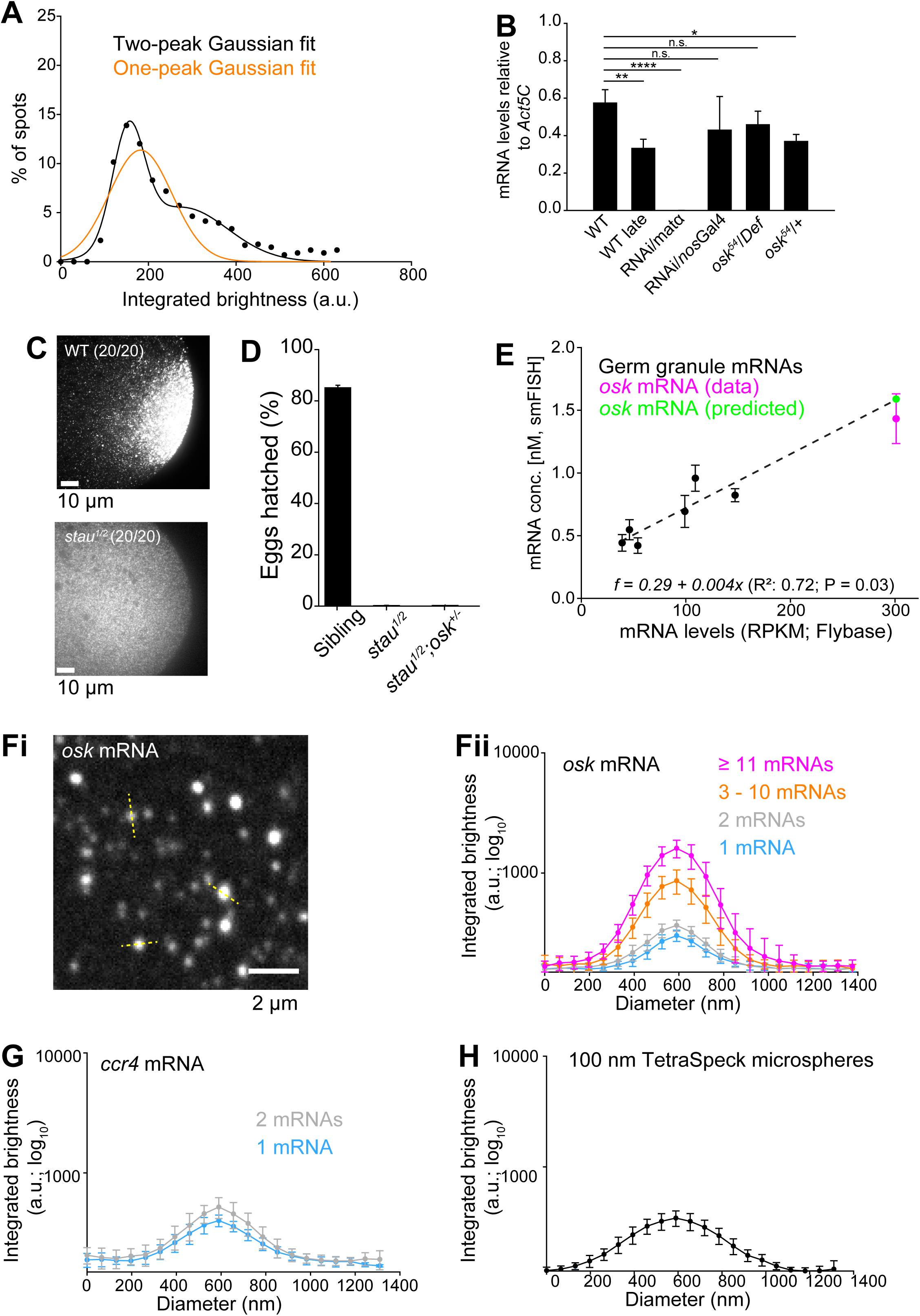
Characterizing mRNA clustering of *osk* in embryos using smFISH. **A** Fitting the distribution of *osk* fluorescent puncta to a two-peak (black) and a single-peak (orange) Gaussian curve for WT embryos. Significant deviation of the fit from the data is observed for the single-peak Gaussian fit (see also Suppl. Table 1). **B** mRNA expression levels of *osk* mRNA determined for WT, WT late, RNAi/*matα*, RNAi/*nosp Gal4*, *osk*^54^*/Df* and *osk*^54^*/+* embryos using qRT-PCR analysis. Mean±STDEV mRNA levels from three biological replicates are shown. *osk* mRNA levels were normalized to *GAPDH2*. Statistical significance: two-tailed *t*-test. *: P = 0.01, **: P = 0.008 and ****: P < 0.001. n.s.: not significant. 3**C** Representative smFISH images of *osk* mRNA at the posterior pole of WT and *stau*^*1*^*/stau*^*2*^ emb3ryos. 20 embryos per genotype were analyzed. **D** Egg hatching rates for *stau*^*1*^*/stau*^*2*^ embryos and their siblings (*stau*^*1*^*/cyo* or *stau*^*1*^*/cyo*) derived from the same cross and *stau*^*1*^*/stau*^*2*^*;osk^+/-^* embryos laid by *stau*^*1*^*/stau*^*2*^*;oskDef/+* females. Data represent a mean±STDEV of three biological replicates. Between 123 and 792 sibling, 214 and 414 *stau*^*1*^*/stau*^*2*^, and 268 and 524 *stau*^*1*^*/stau*^*2*^*;osk^+/-^* embryos were laid per apple juice plate. Statistical significance: two-tailed *t*-test. ****: P < 0.001. **E** Validation of the smFISH quantification. *osk* mRNA concentration (in nM) measured by smFISH was extrapolated from a standard curve using its respective expression levels determined by RNA-sequencing data obtained from Flybase^57^. The standard curve was generated by correlating concentrations determined by smFISH and sequencing of six reference mRNAs (*gcl*, *tao1*, *ccr4*, *sra*, *pum* and *eIF4G2*; dashed line R^2^=0.72). Green circle: somatic concentration of 1.59 nM for *osk* mRNA estimated from the standard curve. Magenta circle and error bars: somatic concentration of 1.43±0.2 nM for *osk* measured by smFISH. **Fi,ii** Image of *osk* mRNA clusters in WT embryos (i; taken from Fig. 1Eii). Yellow dashed lines depict lines scans used to determine the dimensions and linear fluorescence intensity profiles (ii) of *osk* mRNA clusters of different sizes. Mean±STDEV of 20 lines scans for each cluster size are plotted. **G** Mean±STDEV of 20 lines scans for ccr4 monomers (blue) and dimers (gray). **H** Mean±STDEV of 20 lines scans for 100 nm TetraSpeck microspheres.

**SUPPLEMENTAL FIGURE 2:**
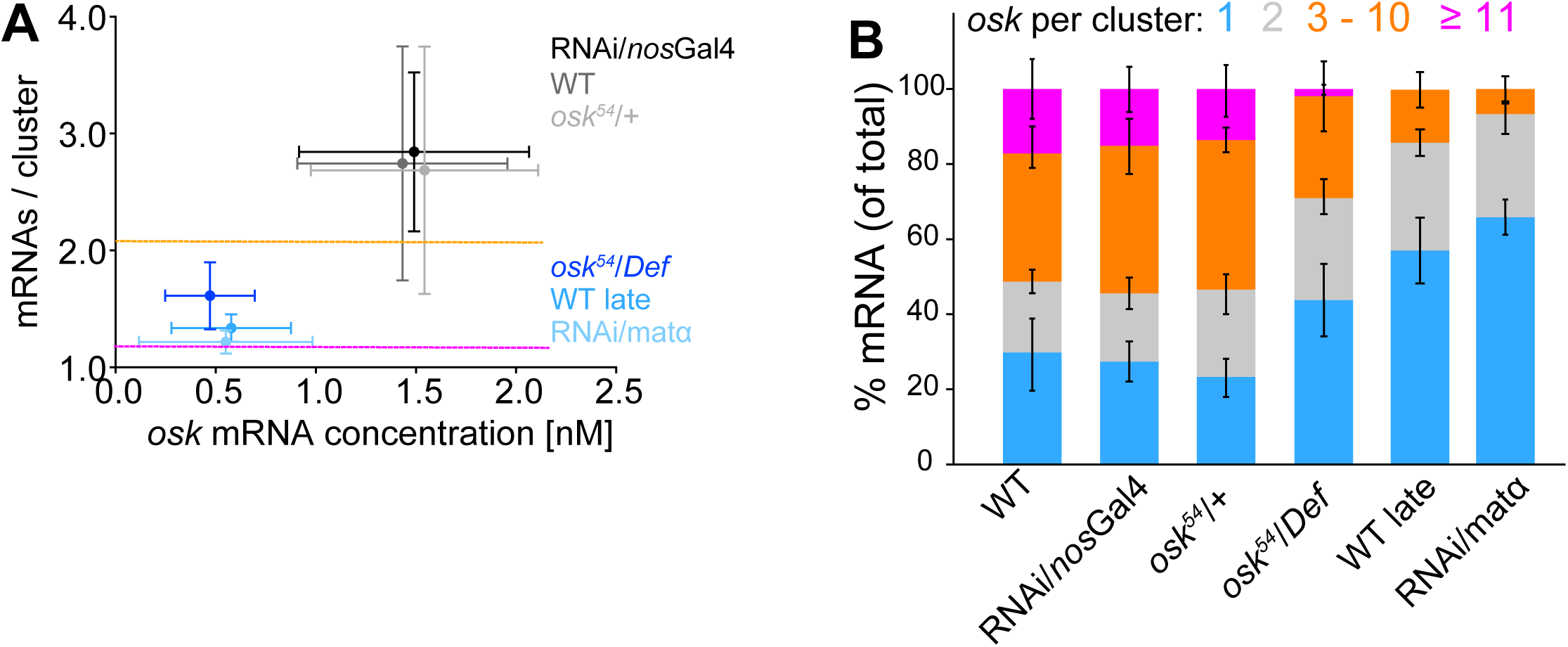
Concentration-driven enhancement of *osk* clustering across different genetic backgrounds. **A** Average *osk* cluster size plotted against the corresponding *osk* mRNA concentration recorded for six different genotypes and developmental stages (Materials and Methods). For comparison, the average (magenta line) ± STDEV (orange line) cluster size of 12 reference mRNAs (*aret, ccr4*, *CG18446*, *eIF4G2*, *gcl*, *orb, Pi3K*, *p53*, *pum*, *shutdown*, *sra* and *tao1*) and their somatic concentrations characterized previously^47^ are shown. **B** Distribution of *osk* mRNA across clusters of varying size. For each cluster size, mean±STDEV from seven WT, WT late, RNAi/*nosp Gal4*, *osk*^54^*/Df* and *osk*^54^*/+* and three RNAi/*matα* embryos is shown.

**SUPPLEMENTAL FIGURE 3:**
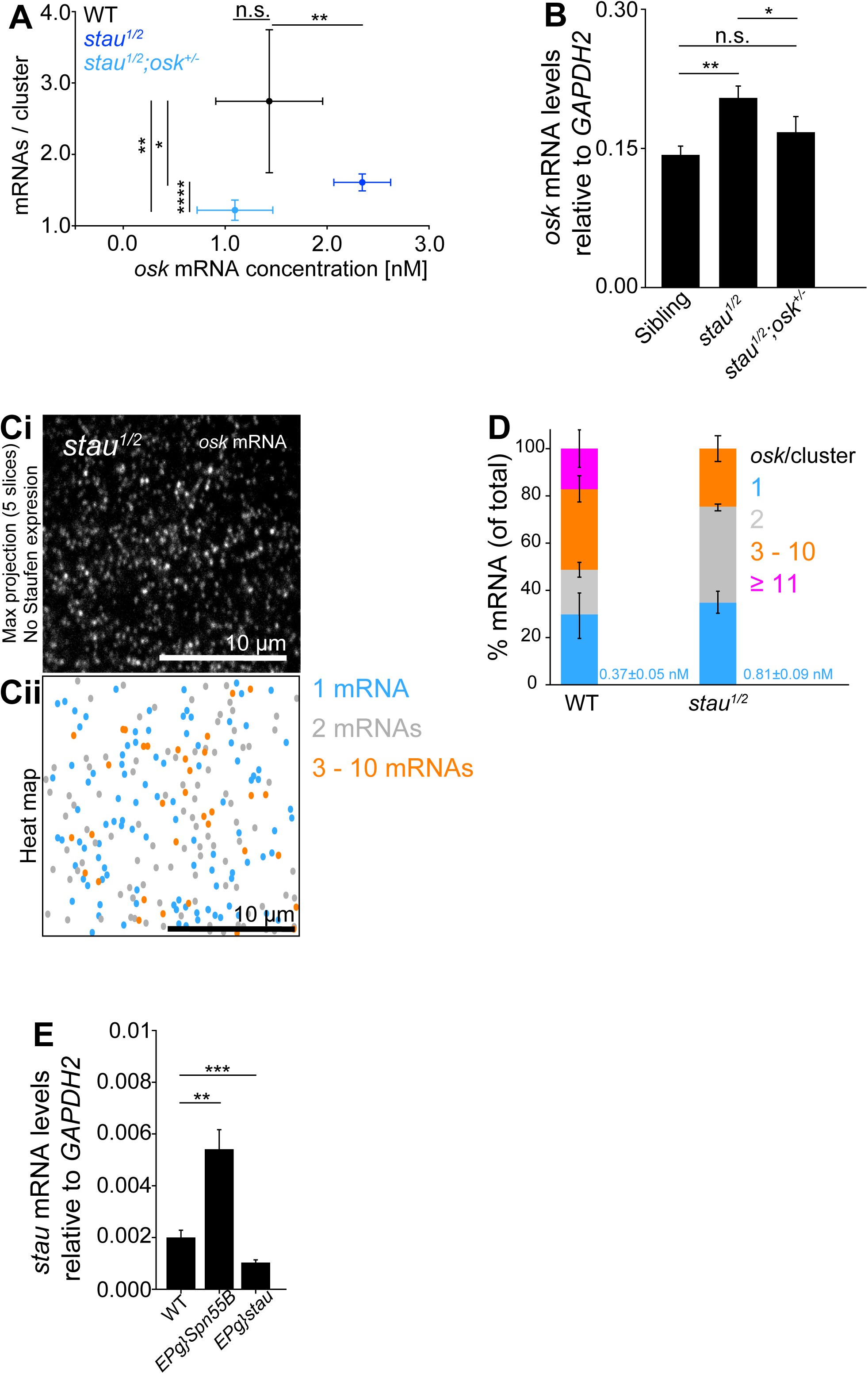
*osk* clustering is diminished in the absence of Staufen expression. **A** Dependence of average *osk* mRNA cluster size on its concentration (nM). Data represent mean±STDEV from seven WT, *stau*^*1*^*/stau*^*2*^ and *stau*^*1*^*/stau*^*2*^*;osk^+/-^* embryos. Data for WT and *stau*^*1*^*/stau*^*2*^*;osk^+/-^* are from Fig. 2F. Statistical significance: two-tailed *t*-test. *: P = 0.01, **: P = 0.002, ****: P < 0.001. n.s.: not significant. **B** *osk* mRNA expression levels determined for WT, *stau*^*1*^*/stau*^*2*^ and *stau*^*1*^*/stau*^*2*^*;osk^+/-^* embryos using qRT-PCR analysis. Mean±STDEV mRNA levels from three biological replicates are shown. *osk* mRNA levels were normalized to *GAPDH2*. Statistical significance: two-tailed *t*-test. *: P = 0.04, **: P = 0.003. **C** *stau* mRNA expression levels determined for WT embryos and those overexpressing *stau* using qRT-PCR analysis. Mean±STDEV mRNA levels from three biological replicates are shown. *stau* mRNA levels were normalized to *GAPDH2*. Statistical significance: two-tailed *t*-test. **: P = 0.007, ***: P = 0.002. n.s.: not significant. **D** Maximal projection of five consecutive optical sections of an ROI, spaced 150 nm apart of *osk* mRNA in *stau*^*1*^*/stau*^*2*^ embryos (i) with a corresponding heat map. **E** Distribution of *osk* mRNA across clusters of varying size. For each cluster category, mean±STDEV of seven embryos per genotype is shown. Data for WT is from Fig. 2G. In blue, the mean±STDEV molarity (n = seven) of *osk* mRNA monomers for each genotype is shown.

**SUPPLEMENTAL FIGURE 4:**
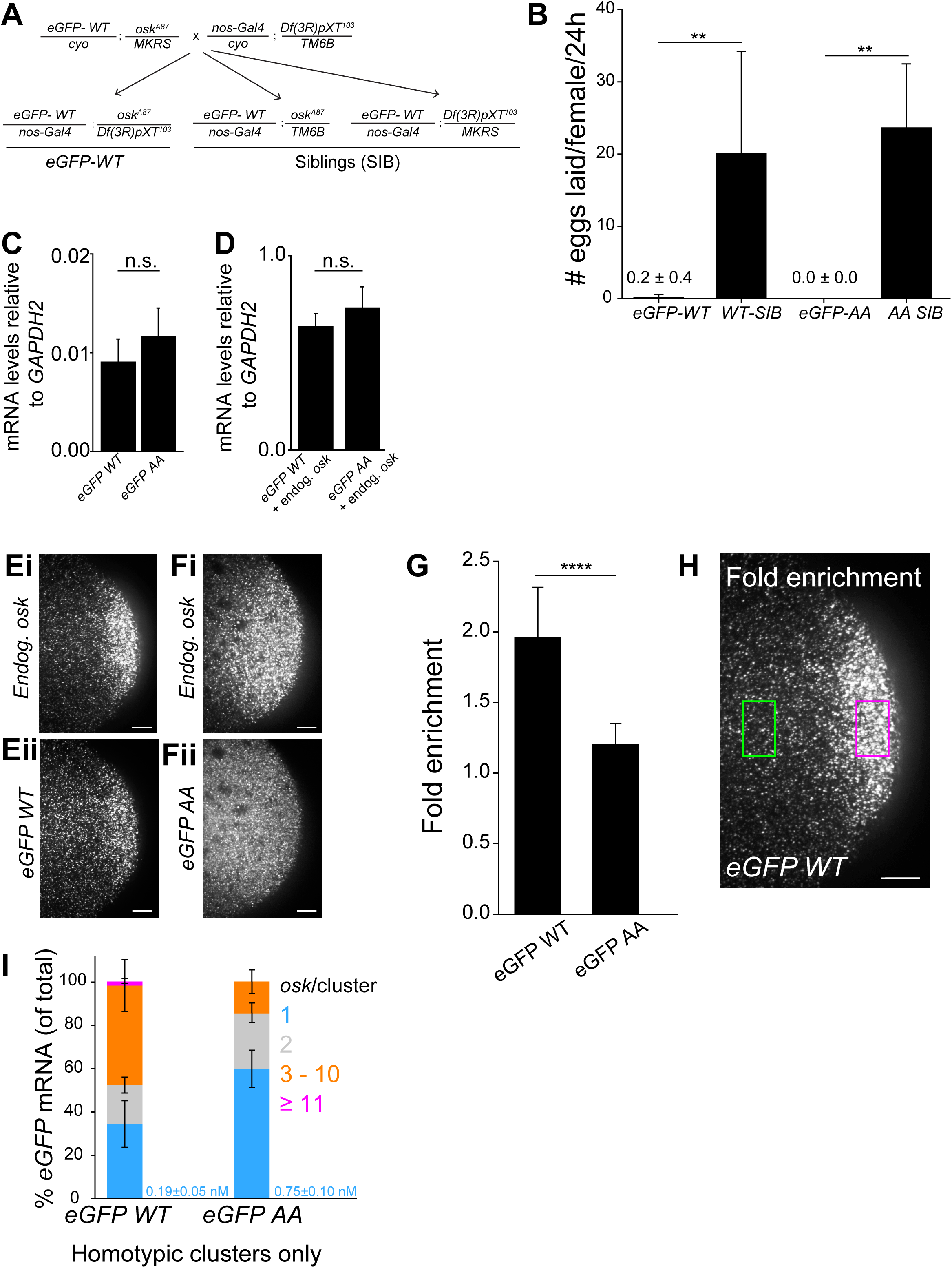
Evaluating expression and localization of eGFP transgenes. **A** Crossing scheme that generated females that expressed *eGFP-WT* or *eGFP-AA* but lacked endogenous *osk* mRNA expression. *osk^A^*^87^ is an RNA null allele of the *osk* gene^59^, while *Df(3R)p^XT^*^*103*^ is an *osk* gene deficiency^49^. **B** The UAS-Gal4/*nanos* promoter system did not drive sufficient expression of *eGFP-WT* and *eGFP-AA* transgenes in *osk^A^*^87^*/Df(3R)p^XT^*^*103*^ females, which lack endogenous *osk* RNA expression, to restore egg laying. Per genotype, 21 three-to-four-day old females were crossed with 10 WT males an allowed to lay eggs on an apple juice plates for 24h (Materials and Methods). Mean±STDEV of six, five, three and four apple juice plates for *eGFP-WT*, *WT-SIB*, *eGFP-AA* and *AA-SIB* females generated in A is shown. Between 0 and 21 eggs were laid per apple juice plate by *eGFP-WT* and *eGFP-AA* females and 176 to 856 eggs were laid per apple juice plate by *WT-SIB* and *AA-SIB* females. Statistical significance: two-tailed *t*-test. **: P = 0.007 and 0.006 for *eGFP-WT* and *WT-SIB* and *eGFP-AA* and *AA-SIB* comparison, respectively. **C,D** mRNA expression levels of *eGFP-WT* and *eGFP-AA* (C) and of *eGFP-WT* and *eGFP-AA* along with the endogenous *osk* mRNA (D). Data represent mean±STDEV mRNA levels from three biological replicates. mRNA levels were normalized to *GAPDH2*. Statistical significance: two-tailed *t*-test. n.s.: not significant. **E,F** Representative smFISH images of embryos expressing endogenous *osk* mRNA (Ei,Fi) and of *eGFP-WT* (Eii) and *eGFP-AA* mRNA (Fii). **G,H** Fold enrichment of *eGFP-WT* and *eGFP-AA* mRNA at the posterior pole (G). Data represent mean±STDEV of 10 (*eGFP-WT*) and 9 (*eGFP-AA*) embryos. Statistical significance: two-tailed *t*-test. ****: P < 0.001. Fold enrichment was calculated by determining the ratio of eGFP fluorescent levels, normalized by the area between the posterior (magenta ROI) and outside (green ROI, Material and Methods) (H). **I** Distribution of *eGFP-WT* and *eGFP-AA* mRNA across clusters of varying sizes. Clusters that co-localized with the endogenous osk were removed from the analysis. Bars represent mean±STDEV cluster size from 13 *eGFP-WT* and 12 *eGFP-AA* embryos. In blue, the mean±STDEV molarity of *eGFP-WT* and *eGFP-AA* mRNA monomers (determined from 13 and 12 embryos, respectively) for each genotype is shown.

**SUPPLEMENTAL FIGURE 5:**
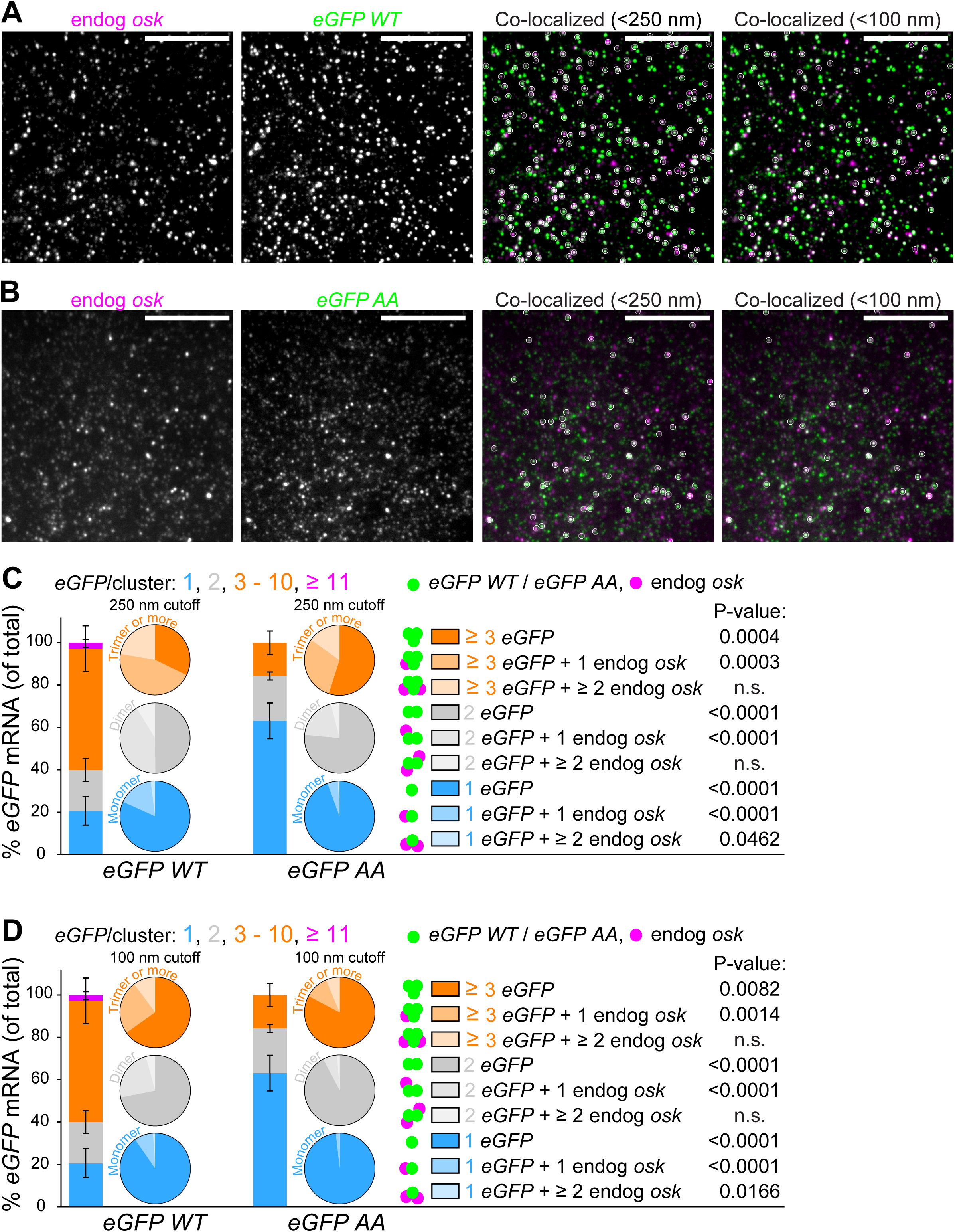
Evaluating co-localization between endogenous *osk* and *eGFP-WT* and *eGFP-AA* mRNA using spot detection analysis. **A,B** Representative smFISH images of co-localizing endogenous *osk* (magenta) and *eGFP-WT* mRNA (A) and *eGFP-AA* mRNA (B) (green). White circles show co-localized *osk* and *eGFP-WT* mRNA or *eGFP-AA* mRNAs at the 250nm and 100 nm cutoff. Scale bar: 10 µm. **C,D** Percent of *eGFP-WT* mRNA and *eGFP-AA* mRNA identified as true monomers, dimers and oligomers using a 250 nm (C) and 100 nm (D) co-localization cut-off. Bars represent mean±STDEV cluster size from 13 *eGFP-WT* and 12 *eGFP-AA* embryos. Pie charts show the composition of each *eGFP* cluster category based on its association with endogenous *osk* mRNA. Differences in heterotypic cluster size between *eGFP-WT* and *eGFP-AA* were evaluated using a two-tailed *t*-test.

**SUPPLEMENTAL FIGURE 6:**
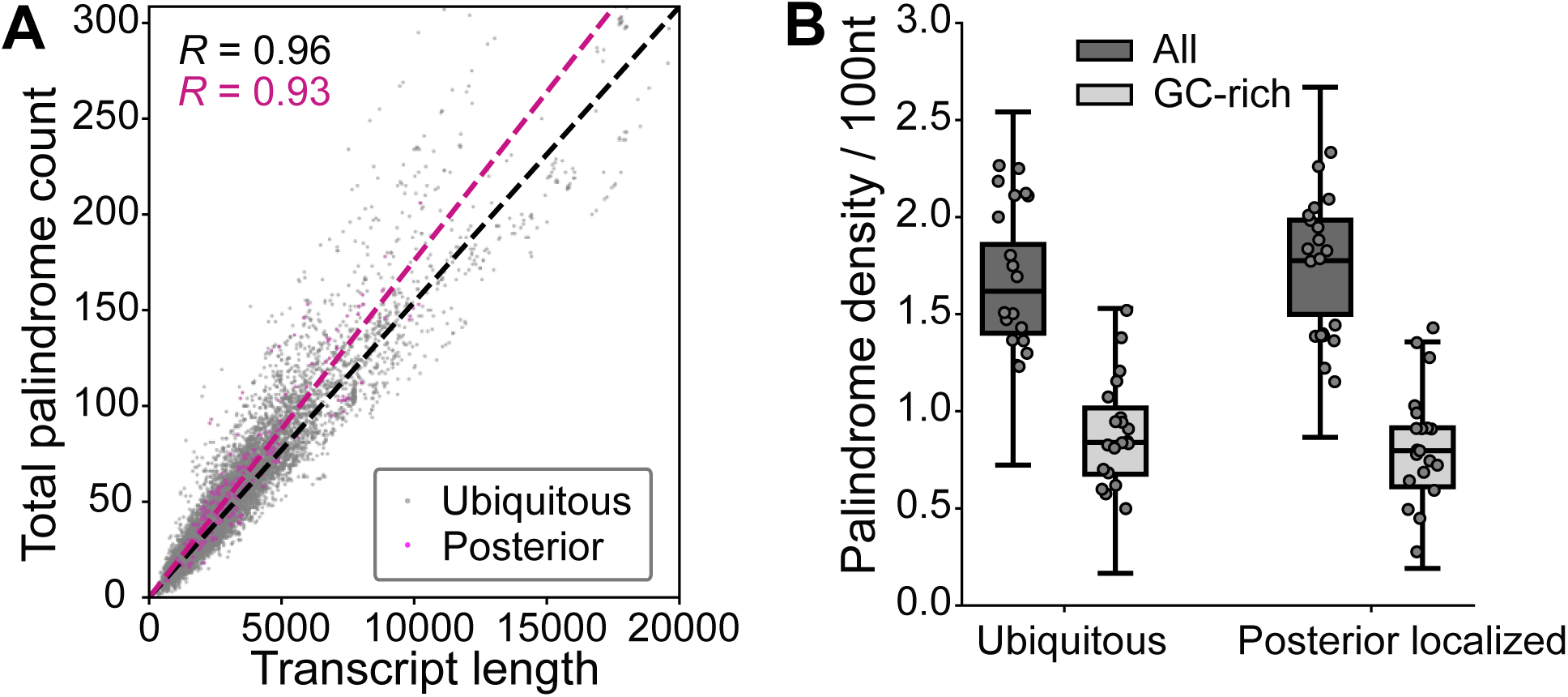
Search for exposed GC-rich palindromes through the early *Drosophila* transcriptome. **A** Scatter plot of the total number of palindromes identified within transcript expressed in the early *Drosophila* embryo as a function of transcript length. **B** Box plot of the palindrome density of ubiquitous and posterior localized *Drosophila* early embryo transcripts. 20 transcripts are selected randomly from each group and overlaid onto the box plot.

## MATERIALS AND METHODS

### Fly lines

Flies were maintained on cornmeal molasses/yeast medium at RT or 25°C using standard procedures^55^. The following fly lines were used: for imaging, embryos expressing Vasa:GFP transgene were used as “wild type” (WT) flies^4^; for egg hatching experiments, *w*^*1118*^ (Bloomington; 5905) were used; *matα-gal4; matα-gal4*^56^; *nosGal4VP16 (w+)/TM3*^88^, *osk* RNAi (y[1] sc[*] v[1] sev[21]; P{y[+t7.7] v[+t1.8]=TRiP.GL01101}attP2; Bloomington; 36903); *osk^Def^* (Df(3R)*pXT103*/*TM3 Sb, Ser*^89^*; yw; osk*^54^ *st ry ss/TM3 Sb*^90,91^; *stau*^*1*^ *cn*^1^ *bw*^1^/*Cyo* (Bloomington; 1507), *cn*^1^ *stau*^*2*^/*Cyo* (Bloomington; 86573)^92^; *osk^A^*^87^/ TM6B (gift from Elizabeth Gavis)^59^; w-; egfp-*oskar* 3’UTR-WT/*cyo*^15^ (termed *eGFP-WT*) and *osk* 3’UTR-AA/*cyo*^15^ (termed *eGFP-AA*), (both gift from Anne Ephrussi’s lab^15^); P{EP}stau[G2356] (w[*]; P{w[+mC]=EP}*stau*[G2356]/CyO, P{ry[+t7.2]=sevRas1.V12}FK1; ; Bloomington; 26989) and P{EPg}*Spn55B*[HP25975] (w[*]; P{w[+mC]=EP}stau[G2356]/CyO, P{ry[+t7.2]=sevRas1.V12}FK1; ; Bloomington; 22094).

### Fly crosses used to modulate *osk* mRNA concentration

To characterize the dependence of *osk* mRNA cluster size on mRNA concentration, we examined embryos laid by females in which *osk* expression was downregulated with Gal4-responsive UAS RNAi using a weak *nanos* (*nos*) promoter^47^ (termed “RNAi/*nosp*”) as well as in embryos laid by *osk*^54^*/Df(3R)p^XT^*^*103*^ (termed *osk*^54^*/Df*) and *osk*^54^*/+* females (Suppl. Fig. 1B, Suppl. Fig. 2A). Here, *osk*^54^ carries a nonsense point mutation which prevents Osk translation^49,50^, triggering nonsense-mediated mRNA decay (NMD)^93^, while *Df(3R)p^XT^*^*103*^ is a deficiency that deletes the *osk* gene^49^. In addition, we also examined embryos in NC 13-14 (termed “WT late”), in which *osk* mRNA levels have diminished due to decay^45^.

We observed that RNAi/*nosp* and *osk*^54^*/+* embryos exhibited similar mRNA expression levels to WT embryos and accordingly formed *osk* mRNA clusters of similar sizes, ranging from 2.69±1.06 (*osk*^54^*/+*) to 2.84±0.68 (RNAi/*nosp*) (Suppl. Fig. 1B, Suppl. Fig. 2A). Additionally, *osk*^54^*/Df* and WT late embryos exhibited similar mRNA expression to RNAi/*matα* and accordingly formed *osk* mRNA clusters with lower sizes ranging from 1.22±0.09 (RNAi/*matα*) to 1.66±0.28 (*osk*^54^*/Df*) (Suppl. Fig. 1B, Suppl. Fig. 2A).

### Egg hatching assays

Approximately 20 *stau*^*1*^*/stau*^*2*^, their sibling *(stau*^*1or2*^*/cyo*), *stau*^*1*^*/stau*^*2*^*;osk^+/-^* and females were crossed with 10 *W*^*1118*^ young males and maintained at 25℃ for three days on standard cornmeal/agar media supplemented with yeast powder. Afterwards, the flies were caged, supplied with an apple juice plate containing a dollop of yeast paste and incubated at 25℃ for 24 hours (hr) to lay eggs^55^. The apple juice plate was then collected, the yeast paste removed and the number of eggs on it determined. The plate was then incubated at 25℃ for an additional 48 hrs, after which the number of hatched eggs counted. The hatching rate was determined by dividing the number of hatched eggs by the total number of eggs on the same plate.

### Egg laying assay

21 *eGFP-WT/nos-Gal4;osk^A^*^87^*/Df(3R)pXT*^*103*^ *(e.i. eGFP-WT), eGFP-AA/nos-Gal4;osk^A^*^87^*/Df(3R)pXT*^*103*^ *(e.i. eGFP-AA), eGFP-WT/nos-Gal4;osk^A^*^87^*/TM6B or eGFP-WT/nos-Gal4; Df(3R)pXT*^*103*^*/MKRS (e.i. WT-SIB)* and *eGFP-AA/nos-Gal4;osk^A^*^87^*/TM6B or eGFP-AA/nos-Gal4; Df(3R)pXT*^*103*^*/MKRS (e.i. AA-SIB)* three-to-four-day old females were crossed with 10 WT males an allowed to lay eggs on an apple juice plates for 24h at 25°C supplemented with a dollop of yeast paste. The apple juice plate was then collected, the yeast paste removed and the number of eggs on it determined.

### Embryo collection

Unless noted otherwise, caged flies were allowed to lay eggs at room temperature (RT) on a fresh apple juice plate containing a dollop of fresh yeast paste for 1.5 hours, after which the embryos were dechorionated, fixed in 4% paraformaldehyde and the vitelline membrane removed by methanol cracking. Embryos were then stored in 100% methanol at 4°C until further use^4,55^.

### Single-molecule fluorescent *in situ* hybridization (smFISH)

48 fluorescently-labeled DNA probes were used to hybridize against *osk*, *Pep* and *eIF3a* mRNA, and 32 were used for eGFP. This approach strongly increases signal-to-noise ration and increased our detection sensitivity, as described previously^4,25,47,55,56^. Subsequent hybridization of mRNAs with smFISH probes was carried out as described before^4,47,55^. smFISH probe sequences against *osk*, *ccr4* and *eGFP* were listed in Trcek, T. et al^4^.

The following 48 smFISH probes against *Pep* were used (listed in the 5′ to 3′ direction):

ccgatccgaacgacgaaaga, taaacactccgcctatccaa, aaacgattctgcggatttcc, gttgacctttgcgttattca, taccacggaaagccatgttg, caaaattgcggttgcggttc, gtccaccatagttgttgttg, caaggcgacatgttcatacc, gtcgcatgttgttaccaaac, gattgttggccaagttgatg, ggttcctgaacagattgttc, aagtcgagaagcgatggtgg, accacgttggttgcgatttc, gtatggagactccttggtag, tagaacatgtcgttcggcac, atgtgcttcttgcacagatg, ccttgatgtggttctcgaaa, cacgcatcatcagatgggtg, aggcgatagctctcctcaat, ctcctgacggatcatgttgg, gatcgacttgagctgctctg, atgcgcttcaagcggtcaaa, gaagttcaggtcgcacatgg, acttgcgatgggtcgagatg, cttcttcagctgcaaatgtc, tgttgcactcaatgcacttg, tcaatgcgagtggcgaactc, cggacagcagatgagtatcg, ctcttcttcggtgctgatgg, tcttgaccggcttcttcttg, tttgtctcgtcagcagcttc, gtcgttgaagtcaaggatgc, cagttgtacttaggtaggcg, ctccagcttggagatcagac, tgcacaccgagcactcatag, acctcggtgtcgaagaactt, cgtacgggagtgaatctcgg, agaagttgcggtgatgagtc, gatcttggtatcgcttgatt, tcaaccttgcgcttcttacg, cgatggatcgtacagctcac, tcagcattatcatccaccat, aggagtttggactggagctg, cggagatggagaggcgtctg, tagtagcgattgtagcgacc, ttacgaattccttcgacgtc, tcctgggcgtgaatgtaaat, atctgttctttcttcatcct.

The following 48 smFISH probes against *eIF3a* were used (listed in the 5′ to 3′ direction):

gttgaggaacattcaagcgc, agattgcgaaagtcctcaga, ttcagtgggttgaaatccac, tagactgaatgcgcttgcac, ccgttctcaatgaagtccac, atgtaaggagtgagcagagc, cattatagtcacgtccttca, cctgagagatctgacgaatc, gcaagagacgctggaactta, atgttgcaaaaactggccag, accagcagtttttccaattc, atctgcatgtcattgtgacg, tgttcttctggtggtcaata, gtcagatcagtgccgaaata, cgagggcatcgattgcaaag, aaccaactgagagcggatct, ccgtgtcagaacagtagaca, ggttcgggtagacgattgaa, aatggtttaccatctggttg, tcgcgatccttgatctcatg, gtcgctgaagaatacgctgg, tttctgtcctcgatgatctt, cacgagcattattctgcttc, ttgcgctctctttcttcttg, gcttggatttcgttctgatg, ttctccttgagactcttctc, cgtctgcgagatttgctgaa, cagcttggaaagcatcttct, tccaacttcttaatgccttc, cgatttgagtttcgactgca, gcgttcgaagtagtcgattt, aaaagcggtatctcctccag, cttttcggccaagtattttt, cagaactccttgtccttaac, aatggcattctcgatgcgtg, ggatacatgcgcttgagacg, aagggcctcaaggaattcat, cttttcgacgtagagggagg, agagcagcctcaaacttttt, aataatgcggtcggctaagc, ctctttctcgcgaaggaagg, tgtgccaagcgaatctcttc, ttttcgtattgggcacgacg, ttccttgattttacgctcag, ttagatgcgcttggacggtc, tgtgaatctggctctcgtcg, cacttgtcatcgcgcgattg, catcaatgcggaaatcgcgg.

Probes were coupled to a fluorescent dye using an in-house labeling protocol as we described previously^25^.

### mRNA fold enrichment

To measure mRNA fold enrichment, defined as the ratio of the mRNA fluorescence between the posterior and outside, the total fluorescence intensity a 3D ROI cropped from the two respective location was determined. The ROI were of the same dimensions. The analysis was performed using ImageJ, as we have done previously^25^.

### Microscopy

Images were acquired in three dimensions (3D) with a 150nm Z step with a vt-instant Structured Illumination Microscope (vt-iSIM; BioVision Technologies). The microscope was equipped with the 405nm 100mW, 488nm 150mW, 561nm 150mW, 642nm 100mW, and 445nm 75 mW lasers, two ORCA-Fusion sCMOS cameras and the Leica HC PL APO 63x/1.30 GLYC CORR CS2, HC PL APO 63x/1.40 OIL CS2 and HC PL APO 100x/1.47 OIL CORR TIRF objectives as described before^56^.

### Line scan analysis to determine the dimensions and fluorescence intensity profiles

Lines scans were performed as described previously^4^. Briefly, per each cluster size, mRNA or bead, 20 lines scans were obtained in Image J and mean±STDEV plotted. 100 nm TetraSpeck microspheres (Thermo Scientific, T7279) were used.

### Quantifying nearest cluster-to-cluster distance

To quantify local *osk* WT and *ccr4* mRNA density, we performed a nearest-neighbor distance analysis on detected cluster coordinates. For each mRNA cluster, the Euclidean distance to its closest neighboring cluster was calculated in three dimensions. Coordinates were converted from pixel units to nanometers using voxel dimensions of 64.5 nm in x and y and 150 nm in z prior to distance computation. Nearest-neighbor distances were computed independently for each cluster without enforcing one-to-one assignment, allowing the same neighboring cluster to be identified for multiple reference clusters where applicable.

### mRNA concentration and cluster size measurements using smFISH

Somatic mRNA concentrations were quantified as described previously^4^. In short, a 3D region of interest (ROI) with known spatial dimensions was cropped from the image and the absolute number of smFISH-labeled mRNAs within the ROI determined using Airlocalize^55,94^, allowing calculation of the nanomolar (nM) mRNA concentration within the ROI, as we have done previously^4,55^.

To determine the average fluorescence intensity of a single smFISH-labeled *osk* mRNA, we initially focused on data obtained from in WT embryos. We plotted the integrated fluorescence intensities of all detected spots as a histogram and fitted it to a single-peak (1) and two-peak (2) Gaussian curve using Sigmaplot, as we have done previously^47,55^:

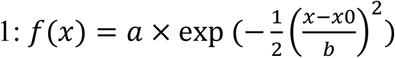

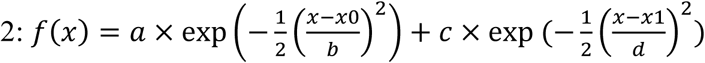

As the fluorescence intensity scales linearly with the number of fluorescently-labeled probes within a diffraction-limited spot^55^, we inferred the mRNA counts by normalizing the cumulative fluorescence intensity of each smFISH-labeled spot against the fluorescence intensity of a single *osk* molecule (e.i. X_0_ value), following established protocols^4,47,55^.

The two-peak Gaussian curve provided a significantly better fit to our data than the single-peak Gaussian model (R^2^ of 0.96 vs. 0.75, respectively; Suppl. Table 1), which displayed a significant deviation from the measured data (Suppl. Fig. 1A). This was anticipated as a considerable proportion of *osk* mRNAs formed clusters. For *osk* mRNA analyzed in WT embryos, this analysis generated mean±STDEV fluorescence intensity of *osk* clusters containing one (X_0_) or two (X_1_) mRNAs (Suppl. Table 1), derived from fits in Suppl. Fig. 1A. Fitting the distribution of *osk* fluorescent puncta for WT embryos to a two-component Gaussian curve revealed two *osk* populations, with mean fluorescence intensities of 154.3±2.9 and 277.2±25.0 arbitrary unites (a.u.), which differed by approximately two-fold (P<0.0001, R^2^:0.96, Suppl. Table 1). We interpreted these as representing clusters containing one and two *osk* mRNAs, respectively. Accordingly, for all subsequent analysis, we fitted the distribution of integrated fluorescence intensities of all detected spots to a two-peak Gaussian model. To estimate an absolute number of mRNAs within each fluorescent spot, we normalized the integrated fluorescence intensity of each smFISH-labeled spot to that of a single *osk* mRNA molecule (X_0_ value), as we have done previously ^4,47,55^.

To determine the size of mRNA clusters (defined as the number of mRNAs per cluster), we analyzed a 3D ROI as described above. Afterwards, we determined the number of mRNAs in each cluster by normalizing its fluorescent intensity to the intensity of a single smFISH-labeled mRNA ^4,55^. Finally, to plot the spatial distribution of *osk* mRNA clusters as a heatmap, we used Sigmaplot as we have done previously^47,55^.

### Quantifying total embryonic mRNA levels using qRT-PCR

Caged flies were provided with fresh apple juice plate with a dollop of fresh yeast paste and allowed to lay eggs for 1.5 hrs, except for the WT late condition, which was aged for an additional hour. After dechorionation, eggs were washed with 1XPBS, and 200 µl TRIzol reagent (Invitrogen; 15596026) was added. Samples were homogenized using a pellet pestle motor (Kimble), followed by addition of 800 µl TRIzol. Samples were stored in -20°C until RNA extraction. Total RNA was purified using the Zymo RNA Clean & Concentrator kit (Zymo; R1018) according to the instructions. cDNA synthesis and qRT-PCR were performed as previously described^25^. *osk*, *stau* and *Gapdh2* primer sequences were described previously^56^. The primer sequences are provided in the Suppl. Table 4. For analysis, *Act5C* or *Gapdh*2 was used as a control to calculate the relative transcript levels, which was 2^−(Cq(gene of interest)−Average of Cq(control))^.

### Co-localization analysis using spot detection

To quantify co-localization between endogenous *osk* mRNA and *eGFP* chimera (WT or AA) transcripts, images were acquired in three dimensions (3D) using an iSIM microscope with a 100x oil objective and subsequently corrected for chromatic aberration using Huygens (Scientific Volume Imaging), as done before^4,47^. smFISH was performed using Quasar 670 labeled probes to detect endogenous *osk* and CalFl590 labeled probes to detect the *eGFP* chimera (WT or AA) transcripts^4^. Spot detection for each fluorescence channel was carried out independently using an algorithm called Airlocalize^4,55^. After thresholding, the algorithm produced 3D coordinates (x, y, z) of all detected spots in pixel units, which were then converted to nanometers based on imaging parameters. Pairwise distances between endogenous *osk* and *eGFP* chimera (WT or AA) transcript spots were calculated using a nearest-neighbor greedy-matching approach, ensuring a one-to-one assignment between spots^4,47^. Co-localized pairs were defined using distance cutoffs of 100 nm and 250 nm. Both cutoffs yielded consistent trends with the 250 nm cutoff data presented in the main text (Fig. 3B-D). Custom Python scripts, mRNA Spot Matcher and mRNA Colocalization Visualizer, were developed to identify and visualize co-localized spots, respectively. Both codes with detailed documentation are available in the X GitHub repository: https://github.com/ayseecer/mRNA_Spot_Matcher_and_Colocalization_Visualizer

### Identification of GC-rich palindromes in the *Drosophila* transcriptome

GC-rich palindromes are identified from fasta sequences downloaded from Flybase using an in-house script. Briefly, the fasta sequences of early Drosophila embryo genes are downloaded from Flybase in bulk, and GC-rich palindromes throughout each fasta sequence are identified as previously described^25^. ViennaRNA RNAfold was used to generate predicted secondary structures of each mRNA sequence. A full list of the identified palindromes is included in Suppl. Table 2, The code for identifying GC-rich palindromes in bulk, and for scoring each palindrome’s intermolecular base pairing propensity (“bp score”), are available in the GitHub repository: https://github.com/AnneyYeZiqing/pfind-bulk-palindrome-finder

